# An integrative oncogene-dependency map identifies unique vulnerabilities of oncogenic EGFR, KRAS, and RIT1 in lung cancer

**DOI:** 10.1101/2020.07.03.187310

**Authors:** Athea Vichas, Naomi T. Nkinsi, Amanda Riley, Phoebe C.R. Parrish, Fujiko Duke, Jenny Chen, Iris Fung, Jacqueline Watson, Matthew Rees, John K. Lee, Federica Piccioni, Emily M. Hatch, Alice H. Berger

**Author notes:** these authors contributed equally to this work (Naomi T. Nkinsi, Amanda Riley). Corresponding author, corresponding author(s): Alice Berger (.

## Abstract

Advances in precision oncology have transformed cancer therapy from broadly-applied cytotoxic therapy to personalized treatments based on each tumor’s unique molecular alterations. Here we investigate the oncogene-specific dependencies conferred by lung cancer driver variants of *KRAS, EGFR*, and *RIT1*. Integrative analysis of genome-wide CRISPR screens in isogenic cell lines identified shared and unique vulnerabilities of each oncogene. The non-identical landscape of dependencies underscores the importance of genotype-guided therapies to maximize tumor responses. Combining genetic screening data with small molecule sensitivity profiling, we identify a unique vulnerability of *RIT1*-mutant cells to loss of spindle assembly checkpoint regulators. This sensitivity may be related to a novel role of RIT1 in mitosis; we find that oncogenic RIT1^M90I^ alters mitotic timing via weakening of the spindle assembly checkpoint. In addition, we uncovered a specific cooperation of mutant *RIT1* with loss of Hippo pathway genes. In human lung cancer, *RIT1* mutations and amplifications frequently co-occur with loss of Hippo pathway gene expression. These results provide the first genome-wide atlas of oncogenic *RIT1*-cooperating factors and genetic dependencies and identify components of the RAS pathway, spindle assembly checkpoint, and Hippo/YAP1 network as candidate therapeutic targets in *RIT1*-mutant lung cancer.

## INTRODUCTION

Somatic mutations that activate EGFR/RAS pathway signaling are a hallmark of lung adenocarcinoma, occurring in more than 75% of tumors^1^. Oncogenes in the EGFR/RAS pathway display ‘oncogene addiction’, a tumor-specific reliance on sustained cell signaling for cell survival, and consequently these mutated oncogenes represent powerful drug targets for lung cancer therapy^2^. Several of the mutated genes in this pathway are clinically targeted to improve outcomes for lung cancer patients. For example, somatic mutations in the *EGFR* proto-oncogene underlie sensitivity to EGFR inhibitors erlotinib and osimertinib^3,4^, and chromosomal rearrangements involving *ALK* underlie sensitivity to crizotinib^5^ and other ALK inhibitors.

Genome-wide CRISPR knockout screens are a powerful tool for associating tumor genotype with unique genetic dependencies^6,7^. This approach has garnered particular interest for the study of oncogenes such as KRAS that for many years lacked direct inhibitors^8,9^. CRISPR-based approaches are also effective for mapping the determinants of drug sensitivity and resistance^10,11^. In lung adenocarcinoma, somatic mutations in EGFR/RAS pathway genes are mutually exclusive^1,12^. Further identification of lung cancer oncogenes and associated therapeutic vulnerabilities is needed to provide targeted therapeutic options for patients whose tumors lack these canonical oncogenic mutations.

Recently, we connected RIT1 to the EGFR-RAS pro-tumorigenic signaling network^12–14^. *RIT1* is mutated in 2% of lung adenocarcinomas^12,13^ and amplified in another 14% of tumors^12^. *RIT1* encodes a ubiquitously-expressed RAS-family GTPase protein^15,16^. Mutations in *RIT1* are mutually exclusive with mutations in *KRAS*, *EGFR*, *ALK*, *MET*, and other driver oncogenes in lung adenocarcinoma, suggesting that *RIT1* may drive EGFR/RAS pathway activation in *RIT1*-mutant tumors^12^. Consistent with this idea, mutant *RIT1* can transform NIH3T3 fibroblasts and confer resistance to EGFR inhibition in *EGFR*-mutant lung cancer cells^13,14^. Mutant *RIT1* has been identified in a patient with acquired resistance to ALK inhibition^17^. Somatic *RIT1* mutations also occur in myeloid malignancies^18^, and focal *RIT1* amplifications are observed in uterine carcinosarcoma^19^. Because *RIT1* mutations are mutually exclusive with other driver alterations in lung adenocarcinoma^12^, patients with *RIT1*-mutant lung tumors have limited therapeutic options of standard chemo- and immuno-therapy regimens but no targeted therapies. Beyond cancer, germline mutations in *RIT1* are found in patients with the “RAS-opathy” Noonan syndrome (NS)^20^.

Both RIT1 protein abundance^21^ and GTP-binding^22^ appear to be central to its oncogenic function. *RIT1* mutations disrupt RIT1’s negative regulation by the ubiquitin adaptor protein LZTR1, leading to increased RIT1 protein abundance. Cancer- and NS-associated variants also display decreased GTP hydrolysis, increased nucleotide exchange, or increased GTP binding^21,22^, and the exact contribution of GTP-binding versus protein abundance to RIT1’s oncogenic function remains to be determined. In all diseases, a similar spectrum of *RIT1* missense and in-frame insertion/deletion mutations is observed, with the majority occurring in the switch II domain of the protein. Mutation at methionine 90 (M90I) is recurrent, but the spectrum of mutations is relatively diverse and the majority have been shown to confer the same cellular transformation capability^13^ and loss of LZTR1 binding^21^. Beyond these observations, relatively little is known about the specific mechanism of action of RIT1, how it differs from KRAS, and what proteins are critical to induce its oncogenic function. Further understanding the cellular consequences of oncogenic *RIT1* mutations could open up new strategies for therapeutic intervention in *RIT1*-mutant cancers and Noonan syndrome.

To investigate the structure of the EGFR/RAS/RIT1 signaling network and identify new therapeutic targets in lung cancer, we performed genome-wide CRISPR screens in isogenic PC9 cells. We find that the dependencies identified broadly confirm expected pathway hierarchy but also identify key differences that highlight the importance of genotype-guided treatment stratification. A key difference we identify is in sensitivity to anti-mitotic therapies; *RIT1*- and *KRAS*-mutant cells differ in their sensitivity to Aurora kinase inhibitors due to a novel role of RIT1^M90I^ in the spindle assembly checkpoint. Furthermore, we identify YAP1 activation as a key cooperating event in *RIT1*-mutant lung cancer. In addition, we identify many other candidate *EGFR-*, *KRAS*-, and *RIT1-*dependencies that should be further explored as novel drug targets for lung cancer therapy. This study expands our knowledge of EGFR/RAS signaling in lung cancer and provides the first genome-wide discovery of factors that cooperate with or antagonize oncogenic RIT1.

## RESULTS

### Context-specific differences in RIT1- and KRAS-stimulated ERK activity

Our previous work found that cancer- and Noonan-associated RIT1 variants such as RIT1^M90I^ can transform NIH3T3 fibroblasts to a phenotype reminiscent of RAS-transformed cells^13^. However, when we examined cellular signaling by Western blot in the same cells, we discovered that oncogenic RIT1 variants fail to promote MEK and ERK phosphorylation, whereas both KRAS^G12V^ and HRAS^G12V^ induce marked increases in MEK/ERK phosphorylation (**Fig. 1a**). Prior data in PC6 pheochromocytoma cells showed that RIT1 variants are capable of stimulating MEK/ERK phosphorylation in that cell type^13^, so we examined several other models to see if RIT1^M90I^ function varies in a cell-type dependent manner. Expression of RIT1^M90I^ in two different human lung epithelial cell lines, SALE or AALE^23^, did increase MEK phosphorylation, but the degree of stimulation differed between the two cell lines despite similar RIT1^M90I^ levels (**Supplementary Fig. 1a**), suggesting that context-dependent factors determine RIT1’s ability to stimulate MEK phosphorylation.

**Fig. 1.**
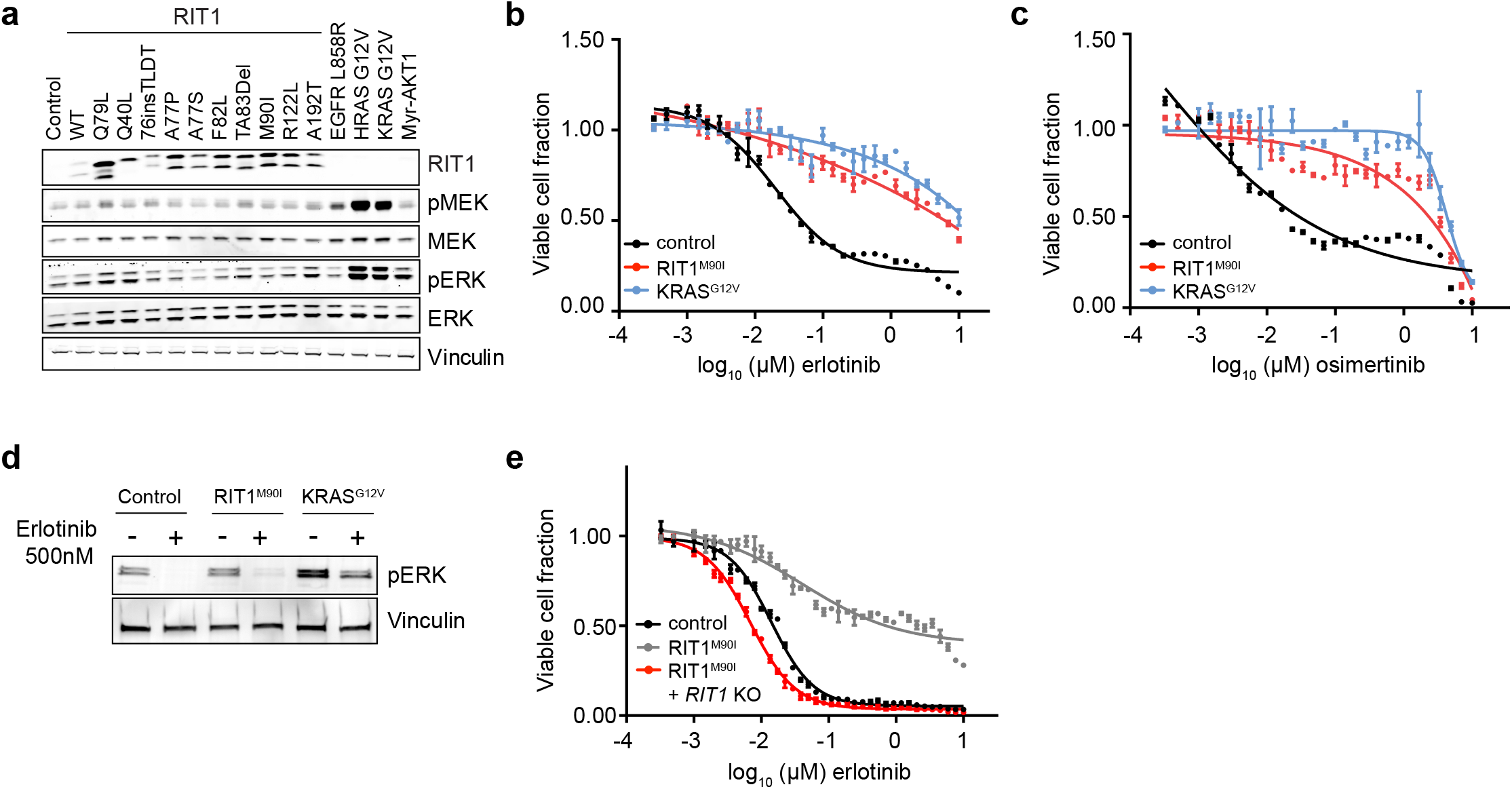
RIT1^M90I^ and KRAS^G12V^ promote resistance to EGFR tyrosine kinase inhibitors. **a**, Western blot of lysates from NIH3T3 cells stably expressing a panel of *RIT1* variants or mutant *RAS*, *EGFR*^*L858R*^, myristolated *AKT1* (Myr-AKT1), or empty vector (control). Vinculin was used as a loading control. p-ERK, phosphorylated Thr202/Tyr204 ERK1/2. p-MEK, phosphorylated Ser217/221 MEK1/2. **b**, Dose-response curve of 96 hours erlotinib treatment in isogenic PC9-Cas9 cells expressing the indicated oncogene or Renilla luciferase (control). CellTiterGlo was used to quantify viable cell number and viable cell fraction determined by normalization to DMSO control. Data shown is the mean ± s.d. of two technical replicates. **c**, Dose-response curve of osimertinib, performed as in (b). **d**, Western blot of lysates from cells shown in (b-c), cultured in the absence or presence of 500 nM erlotinib for 72 hours. p-ERK, phosphorylated Thr202/Tyr204 ERK1/2. Vinculin was used as a loading control. **e**, Dose-response curve of 96 hours erlotinib treatment in clonal *RIT1* knockout (KO) cells compared to parental PC9-Cas9-RIT1^M90I^ cells or control PC9 cells. Data generated and presented as in (b). Unless otherwise indicated, all data are representative results from n = 2 independent experiments.

We previously demonstrated that RIT1^M90I^ and other RIT1 variants, as well as KRAS^G12V^, confer resistance to EGFR targeted therapy in *EGFR*-mutant lung cancer cells^14^. Expression of either RIT1^M90I^ or KRAS^G12V^ in PC9 *EGFR*-mutant lung adenocarcinoma cells renders them resistant to the EGFR inhibitors erlotinib (**Fig. 1b**) or osimertinib (**Fig. 1c**), conferring an almost 1000-fold decrease in drug sensitivity. Despite similar degrees of erlotinib- and osimertinib-resistance in PC9-RIT1^M90I^ and PC9-KRAS^G12V^ cells, induction of ERK phosphorylation in *RIT1*-mutant cells was absent to low, while *KRAS*-mutant cells showed marked rescue of phosphorylated ERK levels (**Fig. 1d**). We confirmed that resistance conferred by RIT1^M90I^ required continued expression of RIT1^M90I^ because erlotinib or osimertinib resistance could be reversed by knockout of *RIT1* using CRISPR-Cas9 technology (**Fig. 1e**, **Supplementary Fig. 1b-d**). These data indicate that stimulation of ERK phosphorylation by RIT1 is cell-type dependent and may involve distinct effectors than KRAS. This finding prompted us to develop a systematic approach to map the differing cellular consequences of oncogenic RIT1 and KRAS activation.

### A genome-scale platform for identification of oncogene-specific dependencies in lung cancer

Genome-scale CRISPR screens have been utilized to identify genetic dependencies of oncogenes in cancer, notably *KRAS*^8,24–26^. Comparative analysis of dependencies in mutant versus wild-type cell lines such as the Broad Institute Dependency Map^6^ are powerful resources that enable the identification of oncogene vulnerabilities. However, the vast majority of mutated cancer genes occur in a relatively low fraction of tumors and cell lines. Indeed, only one lung adenocarcinoma cell line has been identified with a canonical RIT1^M90I^ mutation^13^. An alternative strategy is CRISPR screens of isogenic cell models^8,27^ in which introduced oncogenes confer a “selectable” phenotype to the cells. Because *RIT1* variants can confer resistance to erlotinib in PC9 cells, an *EGFR*-mutant lung cancer cell line^14^, this drug resistance phenotype could provide a powerful screening system to probe the requirements for *RIT1* function in a highly controlled and robust system.

As proof-of-concept, we tested whether small molecule inhibition of downstream signaling components of the EGFR/RAS pathway could overcome RIT1^M90I^-induced erlotinib resistance in PC9 cells. We ectopically expressed RIT1^M90I^ and KRAS^G12V^ in PC9 cells using lentiviral transduction and then cultured the cells in the presence of erlotinib or torin1, a small-molecule inhibitor of mTOR. Co-treatment with torin1 partially re-sensitized cells to erlotinib, demonstrating that both RIT1^M90I^ and KRAS^G12V^ likely act upstream of mTOR to induce erlotinib resistance (**Fig. 2a**).

**Fig. 2.**
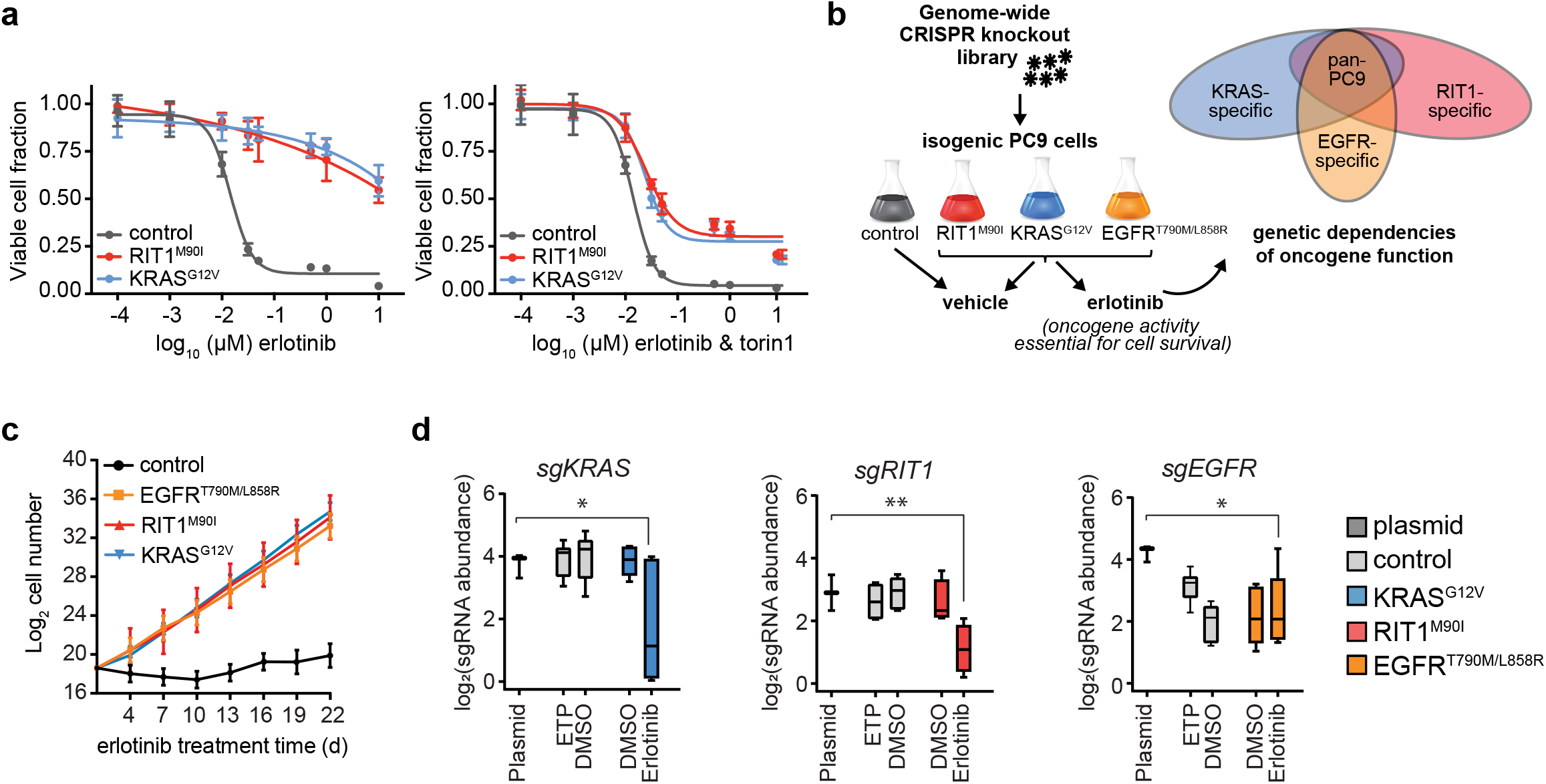
Building an integrative oncogene-dependency map of EGFR^T790M/L858R^, KRAS^G12V^, and RIT1^M90I^. **a**, 96 hour dose-response of isogenic PC9 cells to erlotinib alone (left panel) or to a 1:1 molar ratio of erlotinib and torin1 (right panel). Fraction of viable cells was determined using CellTiterGlo and normalized to the average value in DMSO treated cells. Data shown is the mean ± s.d. of 8 technical replicates. **b**, Schematic of the genome-wide CRISPR-Cas9 screens performed in isogenic PC9 cells. **c**, Proliferation rates of the PC9-Cas9 isogenic cells used for genome-wide screening in 50 nM erlotinib. Data shown is the mean +/− 95% confidence interval for two replicates per cell line. **d**, Box and whiskers plot showing sgRNA abundance (log_2_ reads per million) of sgRNAs targeting each indicated gene. Abundance in the plasmid library, early time point (ETP), or after 12 population doublings in vehicle (DMSO) or erlotinib was determined by PCR and Illumina sequencing. Box plots show the median (center line), first and third quartiles (box edges), and the min and max range (whiskers) of replicates. * *p* < 0.05, ** *p* < 0.01, calculated by MAGeCK analysis (Methods). For control PC9 cells ETP and DMSO n= 3 biological replicates. For oncogene-expressing PC9 cells DMSO and erlotinib n= 2 biological replicates.

To systematically define the factors required for RIT1^M90I^- and KRAS^G12V^-driven drug resistance we developed a pooled genome-wide knockout approach in drug resistant PC9 cells (**Fig. 2b**). Isogenic pools of PC9 cells were generated expressing Cas9 and either a Renilla luciferase vector negative control (“control”), RIT1^M90I^, KRAS^G12V^, or EGFR^T790M/L858R^, an erlotinib-resistant mutant of EGFR used as a positive control (**Supplementary Fig. 2a**). Cas9 activity was confirmed by a sgGFP-GFP flow cytometry assay (**Supplementary Fig. 2b**). Long-term erlotinib resistance was assessed by culturing cells in the presence of 50 nM erlotinib for 21 days (**Fig. 2c**). Whereas cells expressing RIT1^M90I^, KRAS^G12V^, or EGFR^T790M/L858R^ all proliferated at a similar rate in erlotinib, control PC9 cells failed to expand in erlotinib (**Fig. 2c**).

To identify genetic dependencies required for PC9-Cas9-RIT1^M90I^, PC9-Cas9-KRAS^G12V^ and PC9-Cas9-EGFR^T790M/L858R^ cell proliferation in erlotinib, we used the Brunello CRISPR library consisting of 1,000 non-targeting guides and 76,441 guides targeting 19,114 human genes^28^. Cells were transduced, selected with puromycin, and split to erlotinib or vehicle (DMSO) conditions, then maintained in erlotinib or DMSO for approximately 12 population doublings (**Supplementary Fig. 2c**). The change in abundance of each sgRNA was determined by sequencing and comparing the endpoint sgRNA abundance to the initial sgRNA abundance in the plasmid library (**Supplementary Table 1**), which was highly correlated with early time point (ETP) replicates taken just after lentiviral transduction and puromycin selection. Replicate screens were well-correlated (R^2^ range .58 to .94, **Supplementary Fig. 2d**) and exhibited expected lethality of known essential genes with a median strictly standardized mean difference (SSMD) of −3.9 (**Supplementary Fig. 2e, Supplementary Table 1**). To enable quantitative comparisons across screens, we adopted previously established methods^6^ to compute normalized CRISPR scores (CS), scaling the data such that the median CS of all sgRNAs targeting known essential genes is −1 and the median CS of those targeting known non-essential genes is 0 (Methods and **Supplementary Table 2**). As expected, expressed genes had lower CS than non-expressed genes (**Supplementary Fig. 2f**). sgRNAs targeting the expressed oncogene in each cell line were negatively selected in erlotinib-treated cells compared to vehicle-treated cells (**Fig. 2d**), validating the requirement of each oncogene for cell survival.

### No evidence of genetic interaction between KRAS and RIT1

Next we sought to understand whether RIT1 acts upstream, downstream, or in parallel to KRAS. To this end, we evaluated whether knockout of *RIT1* underwent selection in KRAS^G12V^-mutant cells or vice versa. No significant alteration of *RIT1*- or *KRAS*-targeting sgRNAs was observed in the other cell line (**Supplementary Fig. 2g-h**), suggesting that RIT1 and KRAS are not required for drug resistance conferred by the other gene. A similar result was seen for both *RIT1* and *KRAS* sgRNAs in EGFR^T790M/L858R^-expressing cells (**Supplementary Fig. 2i**), suggesting neither *RIT1* nor *KRAS* is required for drug resistance driven by EGFR^T790M/L858R^.

Acquired *KRAS* mutations can drive EGFR inhibitor resistance in patients, but are rarely observed^29^. The low rate of acquired *KRAS* mutations may be due to the mutual exclusivity of *KRAS* and *EGFR* mutations in primary lung adenocarcinoma^12,30^. Work in mouse models has shown that co-expression of mutant *KRAS* and *EGFR* is detrimental to cancer cells, leading to negative selection of cells co-expressing both oncogenes^31^. If true, knockout of *EGFR* should be positively selected in *KRAS*-mutant cells. In support of this model, KRAS^G12V^-mutant cells better tolerated loss of *EGFR*, maintaining high abundance of *EGFR* sgRNAs, in contrast to control PC9 cells or *EGFR-* or *RIT1*-mutant cells (**Supplementary Fig. 2i**).

### Integrative analysis of isogenic CRISPR screen data reveals oncogene-dependent biology

To identify significantly enriched and depleted gene knockouts, we applied MAGECK^32^, which identified an average of 64 positively-selected gene knockouts and 1369 essential genes per cell line (|CS|>0.5 and *p* < 0.05; **Supplementary Table 2**). Of these, 813 were shared across all three isogenic cell lines whereas 238, 209, and 157 were unique to *EGFR*, *KRAS*, and *RIT1*-mutant cells, respectively (**Fig. 3a**, **Supplementary Table 3**). As expected from prior studies^33^, pan-essential genes were significantly enriched for genes in the spliceosome, ribosome, and proteasome pathways (**Fig. 3b**).

**Fig. 3.**
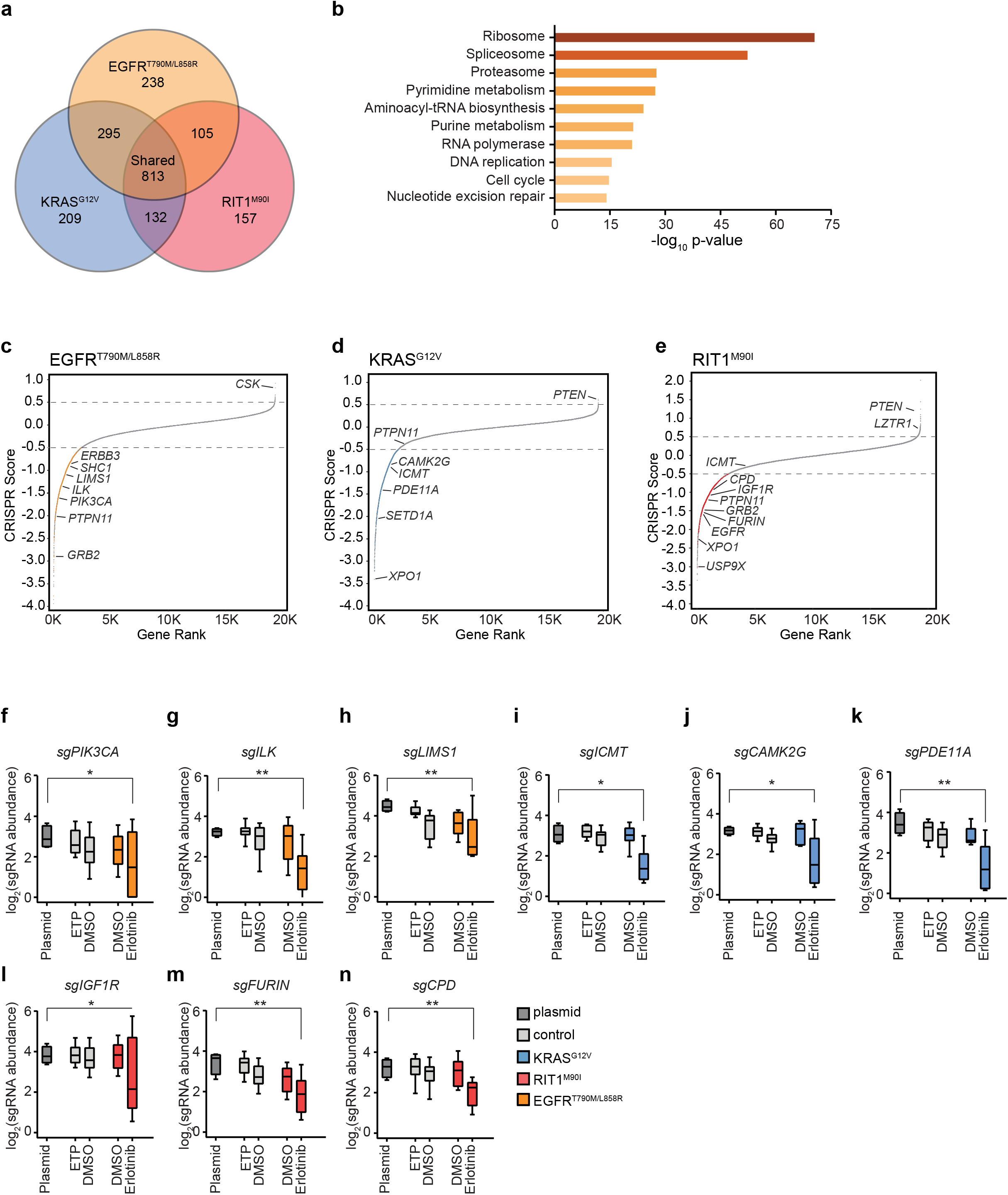
Genome-wide identification of EGFR^T790M/L858R^, KRAS^G12V^ and RIT1^M90I^ oncogene-specific genetic dependencies. **a**, Venn Diagram showing the number of significant essential genes (CRISPR Score < 0.5 and *p* < 0.05) shared or specific to each isogenic PC9 cell line. **b**, MSigDB overlap analysis of the enriched KEGG gene sets in the shared 813 genes from (a). **c-e**, Rank plots of CRISPR scores (CS) of erlotinib-treated vs. starting plasmid in **c**, PC9-Cas9-EGFR^T790M/L858R^ **d**, PC9-Cas9-KRAS^G12V^ and **e**, PC9-Cas9-RIT1^M90I^. Key dependencies discussed in the text are labeled. Gray dashed lines mark genes with |CS| > 0.5. **f-n**, Box and whiskers plot showing the sgRNA abundance (log_2_ reads per million) of sgRNAs targeting each indicated gene. Abundance in the plasmid library, early time point (ETP), or after 12 population doublings in vehicle (DMSO) or erlotinib was determined by PCR and Illumina sequencing. Box plots show the median (center line), first and third quartiles (box edges), and the min and max range (whiskers) of replicates. **f-h**, sgRNAs targeting *PIK3CA*, *ILK*, or *LIMS1* in controls or PC9-Cas9-EGFR^T790M/L858R^ cells. **i-k**, sgRNAs targeting *ICMT*, *CAMK2G*, or *PDE11A* in controls or PC9-Cas9-KRAS^G12V^. **l-n**, sgRNAs targeting *IGF1R*, *FURIN*, or *CPD* in controls or PC9-Cas9-RIT1^M90I^ cells. * *p* < 0.05, ** *p* < 0.01, calculated by MAGeCK analysis (Methods). For control PC9 cells ETP and DMSO n=3 biological replicates. For oncogene-expressing PC9 cells DMSO and erlotinib n=2 biological replicates.

Integration of data from the isogenic cell line screens enabled us to distinguish genes that are broadly essential in PC9 cells from gene knockouts that are negatively and positively selected in cells whose survival is driven by EGFR^T790M/L858R^ (**Fig. 3c**), KRAS^G12V^ (**Fig. 3d**), or RIT1^M90I^ (**Fig. 3e**), respectively. The landscape of these oncogene-specific dependencies differed in specific genes that recapitulated expected biology and pathway hierarchy. For example, in *EGFR*-mutant cells, top essential genes included several known co-receptors and downstream pathway components including; *ERBB3*, *SHC1*, *GRB2*, *PIK3CA*, and *PTPN11*, also known as *SHP2* (**Fig. 3c, 3f**, **Supplementary Fig. 3a-d**). In addition to these well-characterized EGFR downstream components, we found that erlotinib resistance conferred by EGFR^T790M/L858R^ was uniquely dependent on integrin-linked kinase (*ILK*) and its binding partner *LIMS1/PINCH1* (**Fig. 3g-h**). In cancer, ILK and adaptor proteins regulate interactions between tumor cells and the extracellular environment to activate signaling pathways that promote cell proliferation, migration, and epithelial-to-mesenchymal transition^34^. In several human tumors, including non-small cell lung cancer, high ILK and LIMS1 expression correlates with increased disease progression^35–36^ and in *EGFR*-mutant patients, correlates with significantly worse progression-free survival after treatment with EGFR inhibitors^37^. Consistent with a role for ILK in mediating EGFR-TKI resistance, xenograft models of EGFR inhibitor-resistant human hepatocellular carcinoma cell lines found that inhibiting ILK activity increased the sensitivity of cells to EGFR inhibition^38^. Therefore, targeting ILK activity might be a valuable strategy for overcoming EGFR-TKI resistance in patients with EGFR^T790M/L858R^ mutations.

As predicted, *KRAS*-mutant cells were not reliant on upstream RTK signaling genes such as *ERBB3*, *SHC1*, or *GRB2* for survival in erlotinib (**Supplementary Fig. 3a-c**). Although the tyrosine phosphatase gene *PTPN11*, has been recently identified as a potential therapeutic target in *KRAS*-mutant cancers^39^, *PTPN11* knockout did not significantly impact KRAS^G12V^-driven cell survival (**Fig. 3d**, **Supplementary Fig. 3d**). While this difference may be attributed to the different assay systems used, our finding is consistent with earlier reports showing that PTPN11 inhibition is lethal to cells with RTK activation but not to cells with oncogenic RAS proteins^40^. *RIT1-*mutant cells retained sensitivity to *PTPN11* depletion (**Fig. 3e**, **Supplementary Fig. 3d**), highlighting a functional divergence between RIT1^M90I^ and KRAS^G12V^. Instead, KRAS^G12V^-dependencies observed included *ICMT* (**Fig. 3i**), a methyltransferase responsible for the last step in a series of post-translational CAAX-domain modifications required for RAS to associate with the membrane^41^ and recently reported as a *KRAS* dependency^8^. RIT1 lacks a CAAX-domain and as predicted, *RIT1*-mutant cells did not rely on ICMT for survival in erlotinib (**Fig. 3e**). We observed enhanced sensitivity of KRAS^G12V^ cells to the previously reported *KRAS* dependency, *XPO1*^42^. However, loss of *XPO1* was generally lethal to all cell lines screened (**Supplementary Fig. 3e**). Additionally, we identified putative new *KRAS* dependencies including *CAMK2G*, *PDE11A*, and *SETD1A* (**Fig. 3j-k**, **Supplementary Fig. 3f**).

Unlike *KRAS*-mutant cells, RIT1^M90I^-mutant cells were sensitive to knockout of RTK and adaptor protein genes including *PTPN11*, *GRB2*, *SOS1*, and *EGFR* itself. Moreover, RIT1^M90I^-mutant cells required insulin-like growth factor receptor 1 (*IGF1R*) (**Fig. 3l**) and several related components of IGF1R signaling. IGF1R is a multifunctional receptor that promotes cell proliferation, differentiation, and survival through activation of the PI3K-AKT and RAS/MAPK signaling pathways^43^. IGF1R is synthesized as an inactive precursor pro-protein, which requires endoproteolytic cleavage to gain full biological activity^44–45^. In addition to IGF1R, *RIT1*-mutant cells were dependent on the proprotein convertase FURIN (**Fig. 3m**), and carboxypeptidase D (**Fig. 3n**), two proteins directly involved in IGF1R maturation and activity^46–47^.

To validate the genetic dependencies identified, we designed a new custom sgRNA library consisting of 1000 non-targeting sgRNAs and 10,333 new sgRNA sequences targeting 1,288 genes (**Methods, Supplementary Table 4**). We performed secondary screening in three replicates each of vehicle- and erlotinib-treated PC9-RIT1^M90I^ cells. Results from the primary screen and validation screen were highly correlated (R^2^ = 0.77; **Supplementary Fig. 3g**), with 75.4% of essential genes validated and 100% of positively selected genes validating in the secondary screen.

Together, these data illustrate the utility of CRISPR screens in isogenic cell lines for identification of oncogene dependencies in cancer. Comparative analysis of EGFR^T790M/L858R^, KRAS^G12V^, and RIT1^M90I^ dependencies confirmed expectations from established pathway hierarchy. Moreover, the high validation rate in the secondary screen demonstrates the reproducibility of the majority of identified dependencies.

### Enhanced sensitivity of *RIT1*-mutant cells to loss of mitotic regulators

Next, we further investigated the different dependencies conferred by mutant *RIT1* and *KRAS*. To focus on these oncogene-specific dependencies, we calculated a differential CRISPR score between erlotinib- and vehicle-treated screens (**Fig. 4a-b**), and then identified the top differentially essential genes between RIT1^M90I^ and KRAS^G12V^ cells (**Fig. 4c**). *RIT1*-mutant cells retained dependency on several known positive regulators of RAS signaling, including *SOS1* and *SHOC2* (**Fig. 4a,c**), while loss of negative regulators of RAS and genes commonly mutated in Noonan syndrome, such as *NF1*, *SPRED1*, and *LZTR1*, promoted RIT1-induced cell proliferation in erlotinib (**Fig. 4a,c**). In addition to regulators of RAS signaling, several *RIT1* dependencies were involved in mitotic spindle assembly and cell cycle regulation (**Fig. 4a**). The top *RIT1*-specific dependency was *USP9X*, which encodes a deubiquitinase with many target substrates^48–50^. During mitosis, USP9X plays an important role in regulating anaphase initiation and chromosome segregation by stabilizing key mitotic regulators such as Survivin and CDC20^51,48–50^. Knockout of *USP9X* was significantly depleted in RIT1^M90I^ cells in erlotinib, but not in KRAS^G12V^-mutant cells in erlotinib or control PC9 cells in DMSO (**Fig. 4c**). In addition to *USP9X*, *RIT1*-mutant cells were sensitive to depletion of mitotic regulators including *AURKA*, *AURKB*, and *MAD2L1BP*/*p31*^*comet*^ (**Fig. 4c**). Pathway enrichment analysis confirmed that cell cycle and mitotic regulators were significantly depleted in *RIT1*-mutant cells (**Fig. 4d)**. This effect could conceivably be due to differences in proliferation rates, however doubling rates were identical between the two cell lines (**Fig. 2c**). Secondary screening confirmed the enhanced sensitivity of *RIT1*-mutant cells to depletion of *USP9X* and *AURKA* (**Supplementary Fig. 3g, Supplementary Fig. 4a-b**) along with the RAS pathway and Noonan syndrome genes *SHOC2* and *SOS1* (**Supplementary Fig. 3g, Supplementary Fig. 4a-b**).

**Fig. 4.**
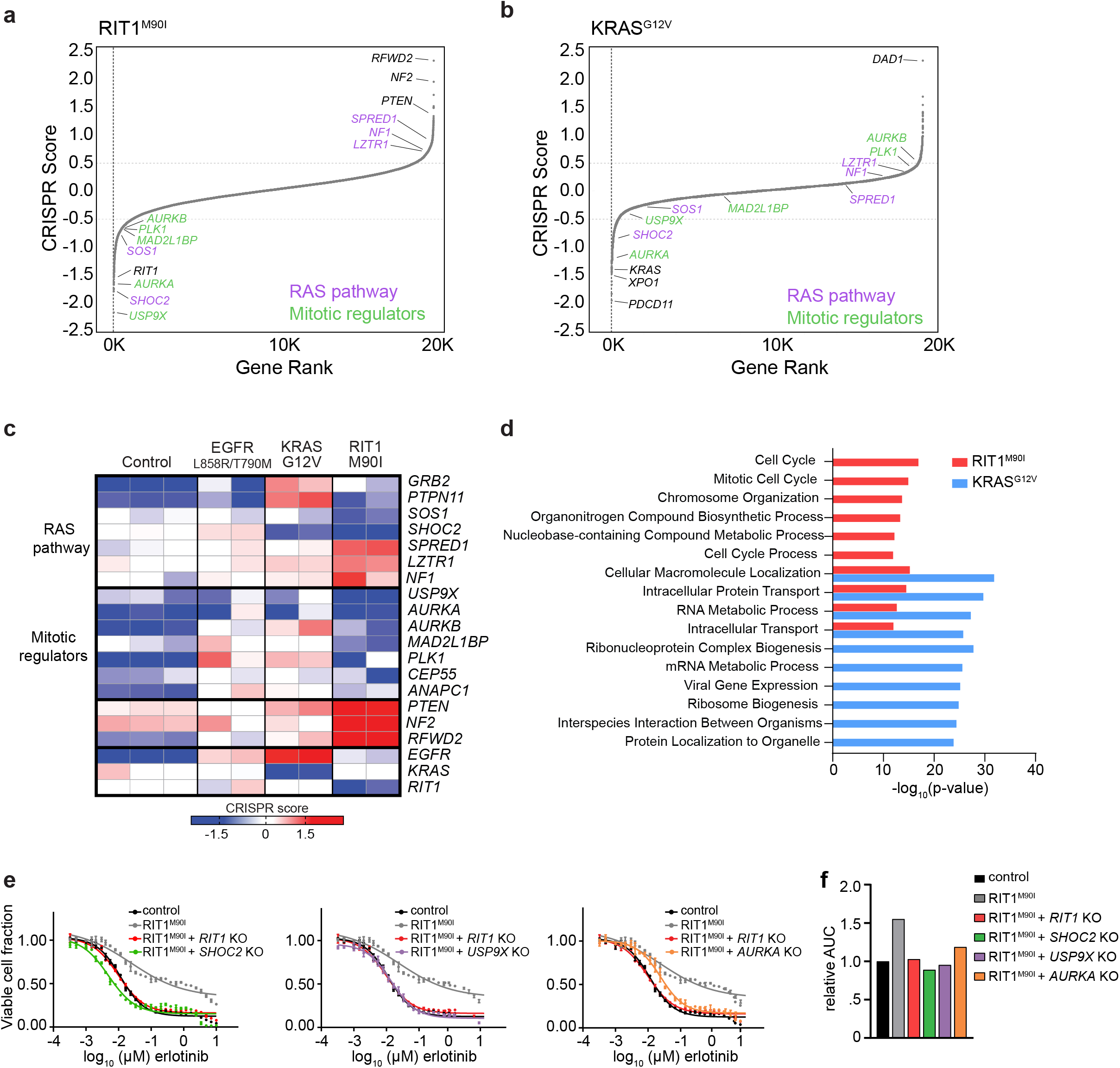
*RIT1*-mutant cells are vulnerable to loss of RAS-pathway and cell cycle genes. **a,b**, Rank plots of erlotinib vs. DMSO differential CRISPR Score (CS) in (a), PC9-Cas9-RIT1^M90I^ or (b), PC9-Cas9-KRAS^G12V^ cells. Key RAS pathway and mitotic regulators are highlighted. Gray dashed lines mark genes with |CS| > 0.5. **c**, Heatmap of CS of selected genes involved in the RAS pathway or mitotic cell cycle regulation clustered by biological pathway. Each column is a different replicate. **d**, MSigDB overlap analysis of the GO Biological Process gene sets significantly enriched in the top 500 PC9-Cas9-RIT1^M90I^ and PC9-Cas9-KRAS^G12V^ essential genes. **e**, 96 hour dose-response curve of erlotinib in clonal *SHOC2* knockout (KO), *USP9X* KO, or *AURKA* KO cells derived from PC9-Cas9-RIT1^M90I^ cells. The same data for control and PC9-Cas9-RIT1^M90I^ cells is plotted on each panel for reference. Data shown is the mean ± s.d. of 2 technical replicates. **f**, Area-under-the-curve (AUC) analysis of data from panel (e).

To further validate the *RIT1*-specific dependency on *USP9X*, *SHOC2* and *AURKA*, we individually knocked out each gene in PC9-RIT1^M90I^ cells. Consistent with the pooled CRISPR screen results, individual loss of *USP9X*, *SHOC2*, or *AURKA* each resulted in restored sensitivity to erlotinib (**Fig. 4e,f**) and osimertinib (**Supplementary Fig. 4e-f)**. These data point to a specific vulnerability of RIT1^M90I^-mutant cells to loss of mitotic regulators, particularly those involved in the spindle assembly checkpoint.

### RIT1^M90I^ weakens the spindle assembly checkpoint

Given that multiple mitotic cell cycle genes were top hits in *RIT1*-mutant cells, we hypothesized that the sensitivity of *RIT1*-mutant cells to genetic loss of spindle assembly checkpoint regulators could correspond to enhanced sensitivity to anti-mitotic therapies. To test this hypothesis, we performed a small molecule screen using 160 clinically-relevant inhibitors in the RIT1^M90I^- and KRAS^G12V^-mutant PC9 erlotinib resistance assay. The majority of compounds affected PC9-RIT1^M90I^ and PC9-KRAS^G12V^ cells with similar efficacy (R^2^ = 0.83; **Fig. 5a and Supplementary Table 6**). Comparison of the area under the curve (AUC) for each compound’s dose-response in each cell line (**Fig. 5b**) revealed that *RIT1*- and *KRAS*-mutant cells shared sensitivity to MEK inhibitors selumetinib and trametinib while also sharing resistance to alkylating agents such as temozolomide and cyclophosphamide (**Fig. 5c and Supplementary Table 6**). A few compounds showed modest selectivity for *KRAS*-mutant cells, including 3 of the 7 RAF inhibitors included in the screen: sorafenib, RAF265, and GDC-0879, whereas top differentially sensitive compounds in PC9-RIT1^M90I^ cells were the Aurora kinase inhibitors alisertib, an inhibitor of Aurora A, and barasertib, an inhibitor of Aurora B (**Fig. 5c**). A PLK1 inhibitor, HMN-214 was also more effective in PC9-RIT1^M90I^ cells. Interestingly, Aurora A, Aurora B, and PLK1 are all important mitotic regulators with multifaceted functions including regulation of the spindle assembly checkpoint^52^. We validated these findings in the same drug resistance assay (**Fig. 5d**) and also sought to determine whether sensitivity to Aurora inhibition was a general phenomenon of RIT1^M90I^-mutant cells. To this end, we tested whether Aurora A/B activity is necessary for *RIT1*-mediated cell transformation in NIH3T3 cells. We found that both alisertib and barasertib treatment suppressed RIT1^M90I^-mediated anchorage-independent growth, whereas transformation by oncogenic RAS was largely unaffected by Aurora inhibition (**Fig. 5e**).

**Fig. 5.**
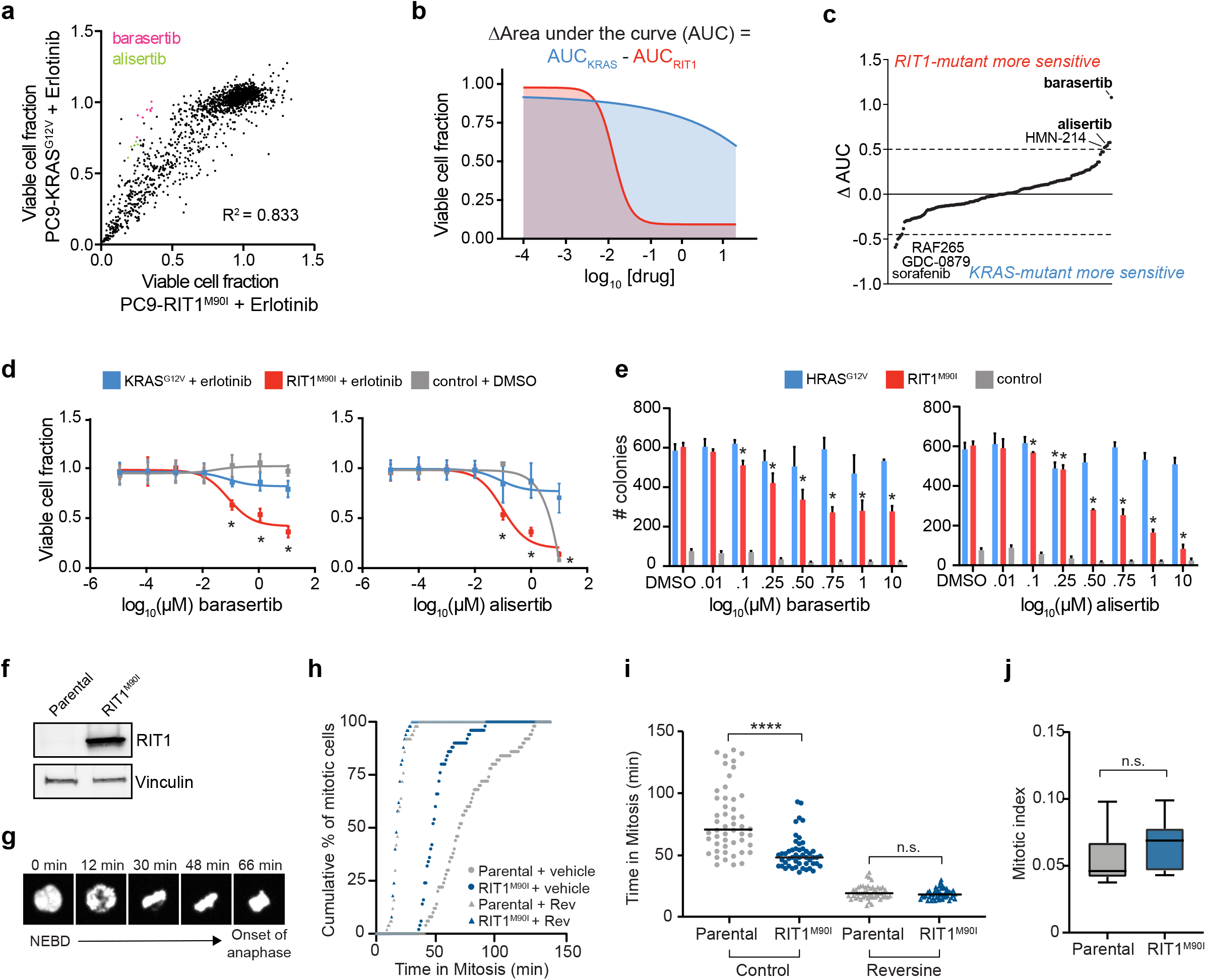
*RIT1*-mutant cells have a weakened spindle assembly checkpoint and are sensitive to Aurora kinase inhibition. **a**, Compound screen of 160 small molecules, 8 doses per compound, in isogenic PC9-RIT1^M90I^ and PC9-KRAS^G12V^ treated in combination with each test condition and 500 nM erlotinib for 96 hours. Data shown is the mean of two replicates of each individual dose/compound. Viable cell fraction was calculated by normalizing CellTiterGlo luminescence in the test condition to CellTiterGlo luminescence of cells treated only with 500 nM erlotinib. **b**, Schematic illustrating differential ‘area-under-the-curve’ (AUC) analysis. **c**,ΔAUC analysis of 160 small molecules generated from data shown in (a) and calculated as in (b). Each dot represents a different compound. ΔAUC are median-centered and plotted from least to greatest. **d**, Validation of enhanced response to barasertib, left, or alisertib, right. Data shown is mean ± s.e.m. of n=16 replicates per dose. * *p* < 0.05 by unpaired two-tailed t-test. **e**, Soft agar colony formation in NIH3T3 cells expressing RIT1^M90I^, HRAS^G12V^ or empty vector (control). Data shown is the mean + s.d. of n=3 replicates. * *p* < 0.05 by unpaired two-tailed t-test. **f**, Western blot of RIT1 expression in parental and RIT1^M90I^-expressing HeLa H2B-GFP cells. Vinculin was used as a loading control. **g**, Duration of mitosis was measured as time from nuclear envelope breakdown to the onset of anaphase. Each frame represents movie stills from time-lapse live-cell imaging of parental HeLa H2B-GFP cells undergoing mitosis. **h**, Time-lapse fluorescence microscopy of time in mitosis in asynchronous parental and RIT1^M90I^-expressing HeLa H2B-GFP cells in normal media conditions (control) or treated with 0.5 μM reversine two hours before imaging. n = 50 cells per condition. Mitotic timing was measured from time of nuclear envelope breakdown to anaphase onset. **i**, Alternative representation of data from (f). Data shown is the mean and individual data points of each condition. **** *p* < 0.0001 by unpaired two-tailed t-test. **j**, Mitotic index calculated as the percentage of mitotic cells in a frame at a chosen time point. Box plots show the median (center line), first and third quartiles (box edges), and the min and max range (whiskers). n = 2 biological replicates, 6 time points per condition, n.s., *p* > 0.05 by unpaired two-tailed t-test.

Fidelity of mitosis relies on the spindle assembly checkpoint (SAC), a critical cellular pathway that senses unaligned kinetochores and arrests mitotic progression during metaphase by inhibiting the anaphase-promoting complex/cyclosome (APC/C) until kinetochores are properly attached to microtubules^52^. Altering the SAC in normal cells accelerates mitotic timing^53^, and in conditions of mitotic stress can either promote mitotic cell death or result in mitotic slippage, the exit from mitosis before proper chromosome alignment is completed^54^. Given that many of the *RIT1* dependencies identified were components of the spindle assembly checkpoint (e.g. USP9X, Aurora kinases, PLK1), we hypothesized that RIT1^M90I^ might weaken the SAC, enhancing the vulnerability of the cells to further loss of SAC activity. To test this hypothesis, we adapted a model system commonly used for mitotic timing experiments^55^, HeLa cells expressing a nuclear H2B-GFP fusion protein^56^. We used live cell fluorescence microscopy to time the duration of mitosis from nuclear envelope breakdown to anaphase onset^53^ (**Fig. 5f,g**). In parental H2B-GFP cells the median duration of mitosis was 70.5 min (95% CI = 63 – 82 min), while in RIT1^M90I^-transduced cells the median duration of mitosis was reduced to 48 min (95% CI = 45 – 51 min) (**Fig. 5h,i**). Overall mitotic index was unaffected, suggesting that mitotic entry is not regulated by RIT1^M90I^ (**Fig. 5j**). The difference in mitotic timing between RIT1^M90I^-expressing cells and parental cells was eliminated by treatment with reversine, an inhibitor of the MPS1 kinase involved in establishing the SAC kinetochore signal, demonstrating that RIT1^M90I^ perturbs mitotic timing at the level of the SAC (**Fig. 5h,i**).

### YAP activation synergizes with RIT1^M90I^ to promote lung cancer

In addition to differences in genetic dependencies, each oncogene displayed substantial differences in the landscape of positively selected gene knockouts (**Fig. 6a-c**). *PTEN* knockout, which is known to promote erlotinib resistance^10,57^, cooperated with both KRAS^G12V^ and RIT1^M90I^ but showed no evidence of selection in EGFR^T790M/L858R^ cells (**Supplementary Fig. 5a**). In contrast, drug resistance driven by EGFR^T790M/L858R^ was promoted by knockout of *CSK* (**Fig. 6a**), which encodes C-terminal SRC kinase, a negative regulator of SRC^58^. SRC activates EGFR via phosphorylation, to enhance EGFR kinase activity and prolong signaling by suppressing EGFR degradation^59^. Enrichment for *CSK* knockout was not observed in *KRAS-* or *RIT1*-mutant PC9 cells (**Fig. 6b-c**). Whereas *EGFR*-mutant and *KRAS*-mutant cells showed few cooperating events (n=37 and n=5 knockouts, respectively), RIT1^M90I^-mutant cells had 152 significantly enriched gene knockouts in erlotinib (**Fig. 6c**). Five of the top 12 genes whose loss synergized with RIT1^M90I^ belong to the Hippo pathway: *NF2*, *CAB39*, *WWC1*, *TAOK2*, and *SAV1* (**Supplementary Fig. 5b-f**). Among all positively-selected gene knockouts (p<0.05, CS>0.5), pathway enrichment analysis using the Molecular Signatures Database (MSigDB) identified the Hippo signaling pathway as the top REACTOME pathway (FDR = 2.08e-5) (**Fig. 6d-e)**. Secondary screen validation confirmed robust enrichment of Hippo pathway knockouts including *NF2*, *CAB39*, and *TAOK1* in RIT1^M90I^-mutant cells (**Supplementary Fig. 3g, Supplementary Fig. 5g**). Validation experiments using pooled and clonal knockout cells demonstrated that loss of *NF2* cooperated with RIT1^M90I^ to promote erlotinib resistance (**Supplementary Fig. 5h-i**).

**Fig. 6.**
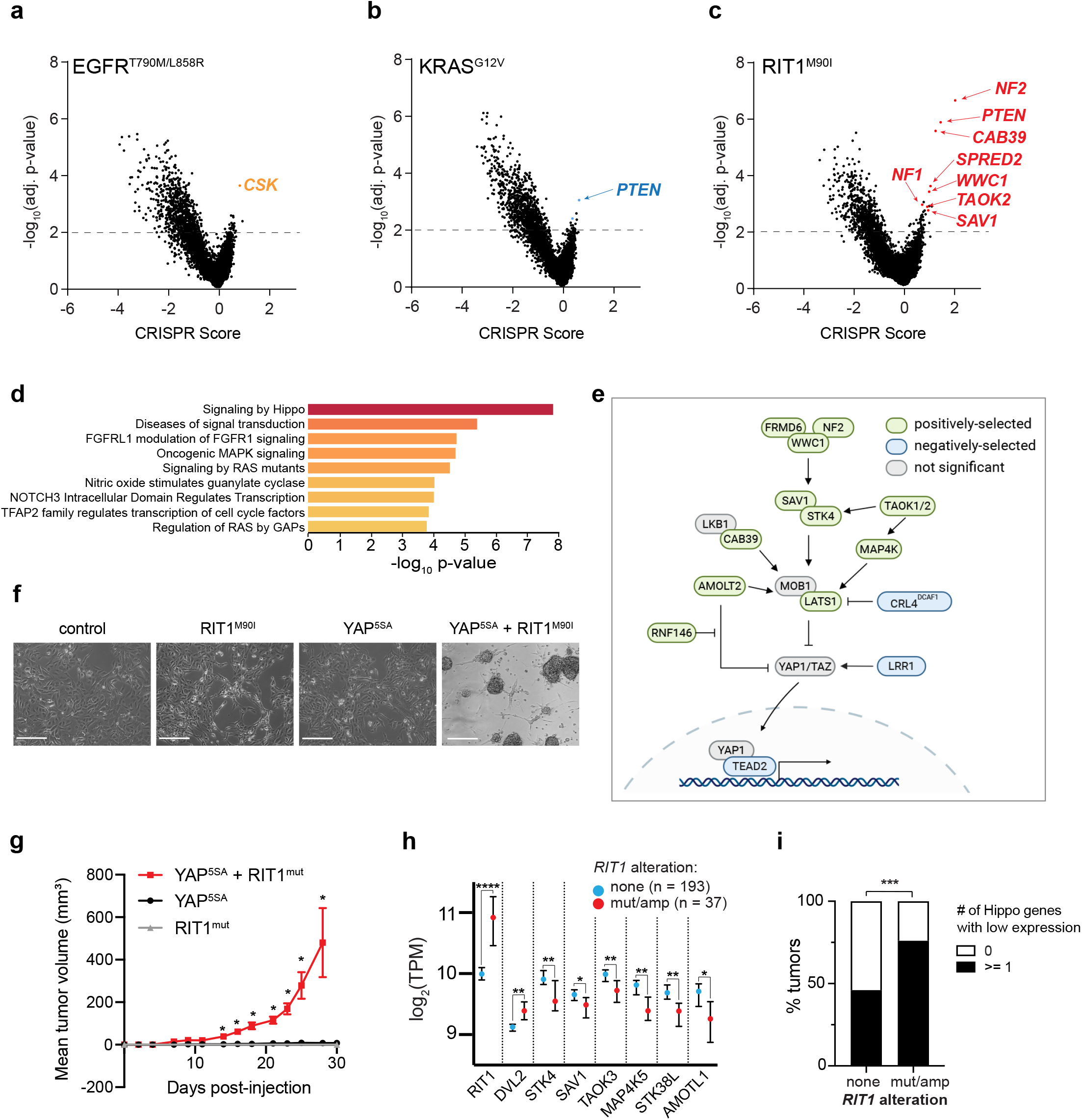
Hippo pathway inactivation synergizes with RIT1^M90I^ to promote cell survival and proliferation. **a-c**, Volcano plots of genome-wide CRISPR screening data from **a**, PC9-Cas9-EGFR^T790M/L858R^ cells, **b**, PC9-Cas9-KRAS^G12V^ cells and **c**, PC9-Cas9-RIT1^M90I^ cells cultured in 40 nM erlotinib for 12 population doublings. “CRISPR Score” indicates the normalized log_2_ (fold-change) of the average of 4 sgRNAs per gene in two biological replicates in erlotinib compared to the starting abundance in the plasmid library. The horizontal dashed line represents the *p*-value threshold of 0.01. **d**, MSigDB overlap analysis of the top REACTOME gene sets in RIT1^M90I^ positively-selected gene knockouts (*p* < 0.05, CRISPR score > 0.5). **e**, Depiction of positively-selected (green) and negatively-selected (blue) Hippo pathway components identified in PC9-Cas9-RIT1^M90I^ (|CS |> 0.5 and *p* < 0.05). **f**, Phase contrast images of SALE-Cas9 cells either alone (control) or expressing RIT1^M90I^, the constitutively nuclear-localized YAP^5SA^, or RIT1^M90I^ and YAP^5SA^. Scale bar = 200 μm. Representative images from *n* = 3 independent experiments are shown. **g**, Xenograft assay of SALE cells expressing RIT1^mut^ (RIT1^T76insTDLT^), YAP1^5SA^, or combined RIT1^mut^ and YAP1^5SA^ in immunocompromised mice. Data shown is the mean + s.e.m. of n=6 tumors per group. * *p* < 0.05 by unpaired two-tailed t-test compared to either YAP1^5SA^ alone. **h**, RNA sequencing data of human lung adenocarcinomas from TCGA. Data shown is the median +/− 95% confidence interval of log_2_-transformed transcripts per million (TPM) in *RIT1*-altered (amplified or mutated, n=37) tumors compared to *RIT1* non-altered tumors (n=193). * *p* < 0.05, ** *p* < 0.01, **** *p* < 0.0001 by unpaired two-tailed t-test. **i**, Proportion of tumors with low expression of any one Hippo pathway gene (Methods). *** p < 0.001 by one-tailed Fisher’s exact test.

The Hippo pathway is a tumor suppressive cellular signaling pathway that regulates proliferation and cell survival via negative regulation of YAP1 stability and nuclear translocation^60^. Enrichment for loss of Hippo pathway genes in RIT1^M90I^ cells suggests that inactivation of Hippo signaling may synergize with oncogenic RIT1 to promote cellular proliferation and survival. However, YAP1 activation has previously been shown to promote erlotinib resistance itself^61^, so we tested whether YAP1 activation can synergize with RIT1^M90I^ in the absence of erlotinib using a human small airway lung epithelial (SALE) cell transformation model^14^. To test whether RIT1^M90I^ or activated YAP1 can transform these cells, we expressed RIT1^M90I^ alone or in combination with YAP1^5SA^, which harbors five serine-to-alanine mutations at critical LATS1/2 phosphorylation sites, resulting in a stabilized, nuclear-localized YAP1 protein^62^. The co-expression of RIT1^M90I^ and YAP1^5SA^ in SALE cells caused the cells to shift from an adherent to a suspension growth phenotype (**Fig. 6f**). Moreover, co-expression of RIT1^mut^ and YAP1^5SA^ synergistically transformed SALE to induce xenograft tumor formation (**Fig. 6g**). Together these data suggest that oncogenic RIT1 and activated YAP1 cooperate to promote tumorigenesis.

To investigate whether co-activation of YAP1 and RIT1 occurs in human lung cancer, we analyzed data from 230 human lung adenocarcinomas sequenced by The Cancer Genome Atlas^12^. Because wild-type *RIT1* overexpression can transform cells^14^ and may confer Noonan syndrome in individuals with germline *LZTR1* mutations^21^, we included *RIT1*-amplified tumors in our analysis. 16% of lung adenocarcinomas harbored mutated or amplified *RIT1* (**Supplementary Fig. 5j**). Tumors showed copy number-related increases in *RIT1* mRNA expression, indicating that *RIT1*-amplified tumors overexpress *RIT1* (**Supplementary Fig. 5k**). Analysis of differentially expressed genes in *RIT1*-mutant and *RIT1*-amplified lung adenocarcinomas identified significant loss of expression of Hippo pathway genes (FDR = 6e-4). Driving this enrichment was the downregulation of key Hippo pathway genes *STK4*, *SAV1*, *TAOK3*, *MAP4K5*, *STK38L*, and *AMOTL1* (**Fig. 6h**). Other significantly altered pathways included EGFR inhibitor resistance (FDR = 2.09E-6), PI3K-AKT (FDR = 2.09E-6), and MAPK signaling (FDR = 7.6E-4) (**Supplementary Table 5**). In contrast, the WNT-pathway gene *DVL2* was overexpressed (**Fig. 6h**), which could further activate YAP1 via crosstalk between WNT and Hippo signaling networks^63,64^. 76% (28/37) of *RIT1*-altered tumors had low expression of at least one Hippo pathway gene, compared to 46% (88/193) of tumors with wild-type/normal *RIT1* (**Fig. 6i**; \*\*\**P* <0.001 by one-sided Fisher’s exact test). Together, these data show that Hippo inactivation synergizes with mutant *RIT1* in cancer models, and Hippo pathway inactivation also occurs in *RIT1*-altered human lung tumors.

## DISCUSSION

Activation of the RTK-RAS signaling pathway is a near-universal hallmark of lung adenocarcinoma^1^, and drug-targetable mutations identified within the pathway have been harnessed to develop genotype-guided therapies for the treatment of lung cancer. While genotype-targeted therapies sometimes result in exceptional tumor responses and have improved lung adenocarcinoma patient survival, selection for tumor cell adaptation occurs rapidly, leading to acquired resistance and necessitating treatment with second- and third-line targeted agents until eventual treatment failure^65^. Highlighting the key requirement for activation of RTK-RAS signaling, acquired resistance usually occurs via second-site mutations in the original oncogene or bypass activation of alternative signaling molecules in the network. Although each mutated oncogene in this network shares the ability to activate this pro-tumorigenic cellular signaling pathway, the overlapping but distinct signaling mechanisms of each protein create unique cellular vulnerabilities.

The role of *RIT1* activation/mutation is a newer and less well-understood part of the RTK-RAS signaling pathway in lung cancer. In this work, we used genetic dependency mapping to uncover the shared and distinct genetic dependencies of oncogenic variants of EGFR, KRAS, and RIT^1^. An advantage of our experimental model system was the specific reliance of the cells on the activity of the expressed oncogene in erlotinib, which presents an advantage over isogenic systems where the phenotype screened has not been firmly linked to the introduced mutation. By intersecting the common dependencies across isogenic cell lines, we were able to exclude pan-essential genes and genes that generally alter sensitivity/resistance to EGFR inhibition, an approach similar to the “Daisy Model” of gene essentiality^66^. This strategy allowed us to overcome the challenging lack of patient-derived experimental model systems in which to study mutant RIT1 function.

RIT1’s involvement in Noonan syndrome and lung adenocarcinoma has confirmed its role in the RAS signaling pathway, although the specific mechanism of its function in this pathway has remained elusive. Although RIT1 is reported to physically interact with C-Raf^21^, its ability to activate ERK appears limited compared to RAS and is cell-type dependent (**Fig. 1a,d** and ref.^21^). In contrast, oncogenic RAS has been studied for decades and is thought to act downstream of the SHP2/PTPN11 phosphatase and SOS1 guanine nucleotide exchange factor to activate effectors including RAF, PI3K, and RalGDS^67^. In agreement with this model, our data show that RAS-mutant cells are insensitive to knockout of *SOS1* and *PTPN11* (**Fig. 4c**). However, other studies suggest that inhibition of PTPN11 can suppress RAS-driven tumorigenesis^39,68^, so further research is required to understand the cell types and contexts where PTPN11 inhibition may be effective. Surprisingly, we find that RIT1^M90I^ differs from KRAS^G12V^ in its dependency on these factors as well as on the SHOC2 scaffolding protein involved in RAF activation (**Fig. 4c**). *RIT1*-mutant cells depend on *PTPN11*, *SOS1*, and *SHOC2*, while showing no requirement for *KRAS* itself. Likewise in *KRAS*-mutant cells, *RIT1* is not required, suggesting KRAS and RIT1 play parallel but distinct roles in regulation of this pathway.

An additional difference between *RIT1*- and *KRAS*-mutant cells is the continued reliance of *RIT1*-mutant cells on RTK complex proteins such as GRB2 and IGF1R. Combined with the requirement for *PTPN11* and *SOS1*, we speculate that RIT1^M90I^ may be involved in activation of RTKs themselves, or drive feedback signaling to RTKs. Further studies are needed to clearly define the structure of RIT1’s requirement for these proteins.

Beyond these classic RTK-RAS signaling components, we uncovered a surprising vulnerability of *RIT1*-mutant cells to perturbation of mitotic regulators, particularly components of the spindle assembly checkpoint. We found that *RIT1*-mutant cells showed heightened sensitivity to loss of mitotic regulators such as *AURKA* and *PLK1*, whether by genetic inactivation or small molecule inhibition, despite no differences in cell proliferation or mitotic index. We showed that RIT1^M90I^ weakens the spindle assembly checkpoint, leaving the cells vulnerable to anti-mitotic therapies. Future genotype-directed clinical trials could determine if patients with *RIT1*-mutant tumors would uniquely benefit from treatment with Aurora kinase inhibitors or other modulators of mitosis. Interestingly, Aurora A activation has been found to drive resistance to the third-generation EGFR inhibitor osimertinib^69^. It is possible that RIT1^M90I^ is harnessing this same mechanism to drive erlotinib resistance and cellular transformation.

One of the strongest genetic interactions we identified in our screen was the pronounced synergy between loss of Hippo pathway genes and *RIT1* mutation. The ability of activated YAP1 and RIT1 to cooperatively transform human lung epithelial cells (**Fig. 6g**) suggests that this synergy occurs not only in the context of drug resistance, but more generally in oncogenesis. Our analysis of human tumor data showed that over 75% of *RIT1*-altered lung adenocarcinomas harbor co-occurring loss of expression of at least one Hippo pathway gene. We hypothesize that RIT1 and YAP1 may act in parallel pathways that synergistically transform epithelial cells. If *RIT1*-mutant tumors require this parallel input, targeting YAP/TEAD activity may be an effective strategy for suppressing *RIT1*-mutant tumorigenesis.

More broadly, our work demonstrates the utility of genome-wide CRISPR screens in isogenic cell lines to identify oncogene dependencies and discover therapeutic targets. Isogenic cell systems are particularly valuable to identify critical dependencies of oncogenes that are mutated in <5% of cases and consequently not well-represented in CRISPR screening databases such as DepMap^6,70^. Use of deeply characterized phenotypes that are closely linked to and predictive of oncogene function is essential. These directed approaches will complement analysis of larger cell line panels towards a goal of building genome-scale dependency maps for all human oncogenes.

## Material and Methods

### Cell Lines

PC9 cells were a gift from Dr. Matthew Meyerson (Broad Institute) and cultured in RPMI-1640 (Gibco) supplemented with 10% Fetal Bovine Serum (FBS). NIH3T3 cells were obtained from ATCC (CRL-1658), and HeLa-H2B cells were a gift from Dr. Daphne Avgousti (Fred Hutchinson Cancer Research Center). NIH3T3 and HeLa-H2B cells were cultured in Dulbecco's Modified Eagle's Medium (DMEM, Genesee Scientific) supplemented with 10% FBS (Sigma). AALE and SALE human lung epithelial cells are immortalized with hTERT and the early region of SV40 and were a gift of Dr. William Hahn (Dana-Farber Cancer Institute) and were cultured in small airway epithelial growth media (SAGM) with SAGM supplements and growth factors (PromoCell or Lonza). All cells were maintained at 37°C in 5% CO2 and confirmed mycoplasma-free.

### Lentivirus Production

Lentivirus was produced using standard triple transfection protocols in HEK293T cells. Briefly, HEK293T were co-transfected with lenti-vector (pLKO/PLX303/PLX317), pCMV-VSV-G (Addgene no. 8454) and psPAX2 (Addgene no. 12260) in FuGENE (Promega) and OptiMEM (Thermo Fisher Scientific). After 48 hours, the supernatant was collected and filtered through a 0.45 μm strainer prior to use or storage.

### Vector construction and cell line generation

For NIH3T3 signaling analysis and soft agar assays, the following plasmids were obtained from addgene; pDONR223-HRAS^G12V^ (addgene no. 82090), pDONR223-EGFR^L858R^ (addgene no. 82906), pDONR223-KRAS^G12V^ (addgene no. 81665), and pBabe-puro-Myr-FLAG-AKT1 (addgene no. 15294). pBabe-puro *RIT1* plasmids have been previously described^6^. Isogenic NIH3T3 were generated by retroviral transduction and selection with 2 μg/ml puromycin. Western blot was used to confirm protein expression.

For cell signaling analysis in SALE and AALE cells, pDONR223-KRAS^WT^ (addgene no. 81751), pDONR223-KRAS^G12V^, and pDONR223-RIT1^M90I^ cDNAs were recombined into pLX317 lentiviral expression vector and lentivirus was generated as above. SALE and AALE cells were transduced with lentivirus followed by selection with 1.5 μg/ml puromycin. Western analysis blot was used to confirm protein expression.

For whole-geome CRISPR knockout screen, PC9 cells stably expressing Cas9 were generated by transducing cells with Cas9 pXPR_111 lentivirus (Genetic Perturbation Platform, Broad Institute) and selecting with blasticidin for 5–7 d. Cas9 protein expression was determined by Western blot. To determine Cas9 activity, PC9-Cas9 cells were transduced with a pXPR_011-sgEGFP (Addgene no. 59702), a vector encoding both GFP and a sgRNA targeting GFP. Following selection with puromycin cells were expanded for 3 days. In parallel, untransfected parental PC9 cells and parental PC9 cells transfected with only pXPR_011-sgEGFP were maintained and used as controls for non-GFP expressing and GFP expressing cells, respectively. GFP expression was analyzed in all three cell lines by flow cytometry and data was analyzed using FlowJo software (Tree Star Inc, Stanford).

To generate isogenic PC9-Cas9 cell lines, pDONR233 vectors encoding EGFR^T790M/L858R^ (addgene no. 82914), KRAS^G12V^ (addgene no. 81665), RIT1^M90I^ (described above), and Renilla Luciferase (addgene no. 25894) were subcloned into the pLX317 lentiviral expression vector. Following verification by Sanger sequencing and restriction digest, stable PC9-Cas9 isogenic cell lines were generated by lentiviral transduction in the presence of 2 μg/ml polybrene followed by selection with 250 μg/mL hygromycin for 48-72 hours. Stable lines were expanded and protein expression confirmed by Western blotting.

For the small molecule screen, PC9-RIT1^M90I^ and PC9-KRAS^G12V^ were generated by transducing parental PC9 cells with lentivirus generated from pLX317-RIT1^M90I^ and pLX317-KRAS^G12V^ generated as described above.

For xenograft analysis, pDONR223-RIT1-p.T76insTDLT (addgene no. 82916) was subcloned into pLX317 and YAP^5SA^ (Addgene no. 42562) was subcloned into pLX311. SALE cell lines cells were generated by transduction of lentivirus generated pLX317-puro-RIT1^T76insTDLT^, pLX311-blast-YAP^5SA^ or both pLX317-puro-RIT1^T76insTDLT^ and pLX311-blast-YAP^5SA^ and selected with 1.5 μg/mL puromycin or 2 μg/mL blasticidin.

To generate isogenic SALE-Cas9 cells, parental SALE and SALE-RIT1^M90I^ cells were transduced with Cas9 lentivirus (addgene no. 52962-LV) and selected with blasticidin. YAP^5SA^ cDNA was PCR amplified from pQCXIH-Myc-YAP-5SA (addgene no. 33093) and cloned via Gibson Assembly ^71^ into the lentiviral FU-CGW vector ^72^ which expresses GFP. SALE-Cas9 and SALE-Cas9 RIT1^M90I^ cells were transduced with YAP^5SA^ lentivirus and 48 hours post-transduction GFP-expression was confirmed by direct fluorescent expression using the EVOS FL Color imaging system (Thermo Fisher Scientific).

### Genome-wide CRISPR gene knockout screen

The human CRISPR Brunello lentiviral prep was obtained from the Broad Institute Genetic Perturbation Platform and is also available from Addgene (73179-LV). The library contains 76,441 sgRNAs targeting 19,114 protein-coding genes and 1,000 non-targeting control sgRNAs. For genome-wide CRISPR screening, 320 million PC9-Cas9-Luciferase, PC9-Cas9-RIT1^M90I^, PC9-Cas9-KRAS^G12V^, or PC9-Cas9-EGFR^T790M/L858R^ cells were infected with the Brunello Library lentivirus (Doench et al., 2016) at a low MOI (<0.3). At 24 h after infection, the medium was replaced with fresh media containing 1 μg/mL puromycin (Sigma). After selection on day 7, cells were split into 2 replicates containing 40 million cells each and treated with either DMSO (Sigma-Aldrich) or 40nM erlotinib (Selleckchem). Cells were then passaged every 3 days and maintained at 500-fold coverage. For early time point analysis (day 7) an initial pool of 60 million cells was harvested for genomic DNA extraction from each of the cell lines. After ~12 doublings, a final pool of 60 million cells was harvested in ice-cold PBS and stored at −80°.

Genomic DNA was extracted using the QIAamp DNA Blood Maxi Kit (QIAGEN) and the sgRNAs from each sample were PCR amplified by dividing gDNA into multiple 100 μl reactions containing a maximum of 10 μg gDNA following Broad Institute standard protocols. Per 96-well plate, a master mix consisted of 150 μl ExTaq polymerase (Takara Bio), 1,000 μl of 10x ExTaq buffer (Takara Bio), 800 μl of dNTP (Takara Bio), 50 μl of P5 primer (stock at 100 μM concentration), and 2,075 μl water. Each well consisted of 50 μl gDNA plus water, 40 μl PCR master mix, and 10 μl of P7 primer (stock at 5 μM concentration). PCR cycling conditions: an initial 5 min at 95 °C; followed by 30 s at 95 °C, 30 s at 53 °C, 20 s at 72 °C, for 28 cycles; and a final 10 min extension at 72 °C. PCR samples were purified with Agencourt AMPure XP SPRI beads (Beckman Coulter). Samples were sequenced on a HiSeq 2500 (Illumina). Raw FASTQ files were demultiplexed and sgRNA counts were calculated using PoolQ v2.2.0.

P5/P7 primers were purchased from Integrated DNA Technologies (IDT). [s] = stagger region, [NNNNNNNN]= barcode

P5-5’AATGATACGGCGACCACCGAGATCTACACTCTTTCCCTACACGACGCTC TTCCGATCT[s]TTGTGGAAAGGACGAAACACCG 3’

P7-5’CAAGCAGAAGACGGCATACGAGATNNNNNNNNGTGACTGGAGTTCAGAC GTGTGCTCTTCCGATCTTCTACTATTCTTTCCCCTGCACTGT 3’

### CRISPR Validation Library and Screening

For secondary screening, we generated a custom library containing 1,000 non-targeting control sgRNAs and 11,333 sgRNAs targeting 1,288 protein-coding genes (**Supplementary Table 7**). For validation screening, 180 million PC9-Cas9-RIT1^M90I^ were infected with the validation library lentivirus at a low MOI (<0.3). At 24 h after infection, the media was replaced with fresh media containing 1 μg/mL puromycin (Sigma). After selection on day 7, cells were split into 3 replicates containing 6.6 million cells each and treated with either DMSO (Sigma-Aldrich) or 40nM erlotinib (Selleckchem). Cells were then passaged every 3 days and maintained at 500-fold coverage. For early time point analysis (day 7) an initial pool of 20 million cells were harvested for genomic DNA extraction from each of the cell lines. After ~12 doublings, a final pool of 20 million cells were harvested for genomic DNA extraction using the QIAamp DNA Blood Maxi Kit (QIAGEN). PCR and sequencing were carried out as described above.

### Single gene knockout generation and genotyping

To induce *RIT1*, *NF2*, *USP9X*, *AURKA*, or *SHOC2* gene knock-out in PC9-Cas9-RIT1^M90I^ cells, a vector-free CRISPR-mediated editing approach was used. Briefly, cells were co-transfected using lipofectamine CRISPR max (Life technologies) with three gene-specific synthetic guide RNAs (Synthego, **Supplementary Table 7**). For each single-gene knockout cell line, gene editing was confirmed by Sanger sequencing. Custom oligos flanking the targeted sites were used to amplify genomic DNA from pooled edited cells (**Supplementary Table 7**) using High-Fidelity 2× Master Mix (New England Biolabs). Indel frequencies were quantified by comparing unedited control and knockout cell lines using Inference of CRISPR Edits (ICE)^73^.

### Cell Lysis and Immunoblotting

Whole-cell extracts for immunoblotting were prepared by incubating cells on ice in RTK lysis buffer [20 mM Tris (pH 8.0), 2 mM EDTA (pH 8), 137 mM NaCl, 1% IGEPAL CA-630, 10% Glycerol] plus phosphatase inhibitors (Roche) and protease inhibitors (cOmplete, Mini, EDTA-free, Roche) for 20 minutes. Following centrifugation (>16,000*g* for 15 minutes), protein lysates were quantified using the Pierce BCA Protein Assay Kit (Thermo Fisher Scientific). Lysates were separated by SDS–PAGE and transferred to nitrocellulose or PVDF membranes using the Trans-blot Turbo Transfer System (BioRad) or iBlot (Invitrogen). Membranes were blocked in 1x Tris Buffered Saline (TBS) with 1% Casein (BioRad) for 1 h at room temperature followed by overnight incubation at 4 °C with primary antibodies diluted in blocking buffer. StarBright (BioRad) or IRDye (LiCOR) secondary antibodies were used for detection and were imaged on ChemiDoc MP Imaging System (BioRad). Loading control and experimental protein(s) were probed on the same membrane in all cases. For clarity, loading control is cropped and shown below experimental condition in all panels regardless of the relative molecular weights of the two proteins.

Primary antibodies used for immunoblotting: Phospho-p44/42 MAPK (Erk1/2) (Thr202/Tyr204) (Cell Signaling Technology, 4370), p44/42 MAPK (Erk1/2) (Cell Signaling Technology, 9107), Phospho-MEK1/2 (Ser217/221) (Cell Signaling Technology, 9154), MEK1 (Cell Signaling Technology, 2352), Cas9 (Cell Signaling Technology, 14697), EGFR (Cell Signaling Technology, 2239), β-Actin (Cell Signaling Technology, 4970), Vinculin (Sigma, V9264); Firefly Luciferase (Abcam, Ab16466), RIT1 (Abcam, Ab53720), KRAS (Sigma, WH0003845M1), NF2 (Abcam, Ab109244), USP9X (Proteintech, 55054-1-AP).

### Drug Treatment and Proliferation Analysis

For proliferation assays cells were plated in 384-well plates at a density of 800 cells per well in 40 μl total volume. One day later, a serial dilution of each inhibitor was performed using a D300e dispenser (Tecan). 96 hours post-treatment, cell viability was determined using CellTiterGlo reagent (Promega) and luminescence quantified on an Envision MultiLabel Plate Reader (PerkinElmer). To calculate the fraction cell viability drug-treated cells were normalized to average cell viability of DMSO-only treated cells. Curve fitting was performed using GraphPad Prism four parameter inhibitor response with variable slope. AUC values were calculated by GraphPad Prism (Graphpad). Inhibitors were obtained from SelleckChem: Erlotinib-OSI-744 (S1023), Osimertinib-AZD9291 (S7297), Torin 1 (S2827), Alisertib-MLN8237 (S1133), Barasertib-AZD1152 (S1147). DMSO (Sigma-Aldrich).

### Small Molecule Drug Screen

PC9-RIT1^M90I^ and PC9-KRAS^G12V^ were plated in 384-well plates at a density of 800 cells per well in 40 μl total volume. One day after seeding, cells were treated with 500 nM erlotinib in combination with the small molecule library (a kind gift of Dr. Stuart Schreiber, Broad Institute). Each small molecule was tested across an 8-point dilution series (**Supplementary Table 6**). 96 hours post-treatment cell viability was determined using CellTiterGlo reagent (Promega) and luminescence quantified on an Envision MultiLabel Plate Reader (PerkinElmer). To calculate the fraction cell viability drug-treated cells were normalized to 500 nM erlotinib only treated cells. Dose–response curves were plotted using GraphPad Prism software (GraphPad); AUC values were generated using GraphPad Prism software and three parameter inhibitor response setting. Delta-AUC was calculated by subtracting the AUC for each compound in RIT1^M90I^ from the AUC of the same compound in KRAS^G12V^ cells. Each experiment was carried out in two technical replicates.

### Soft Agar Assays

For soft agar assay, 5 × 10^3^ NIH3T3 cells expressing *RIT1^M90I^*, *HRAS^G12V^*, or control (empty vector) were suspended in 1 ml of 0.33% select agar in DMEM/FBS and plated on a bottom layer of 0.5% select agar in DMEM/FBS in six-well dishes. Each cell line was analyzed in triplicate. Colonies were photographed after 14–21 days and quantified using CellProfiler ^74^. For soft agar inhibitor experiments, alisertib (MLN8237) or barasertib (AZD1152) was suspended in the top agar solution at a final concentration of .01-10 μM.

### In vivo xenograft study

Athymic nude (nu/nu) mice (4 to 6 weeks old) for the xenograft study were obtained from (Jackson Laboratory, Bar Harbor, ME, USA). SALE cells expressing *RIT1^T76insTDLT^*, *YAP^5SA^*, or *RIT1^T76insTDLT^* + *YAP^5SA^* were harvested by trypsinization, washed in PBS and resuspended at 10^6^ cells/ml in PBS. Two hundred microliters (2 × 10^6^ cells) were injected into each injection site, n=2 injection sites/mouse; 3 mice/condition for 6 replicates per cell line, in 4–6-week-old nu/nu mice. Cells were allowed to engraft for 1 week, then tumors were measured every 2-3 days using a digital caliper (VWR, Radnor, PA, USA). Tumor volume was calculated with the formula 0.5 × L × W^2^ where L is the longest diameter and W is the diameter perpendicular to L. Tumors were monitored until the largest tumor reached 2 cm in diameter. All animal experiments were carried out in accordance with the Dana-Farber Cancer Institute Institutional Animal Care and Use Committee guidelines.

### Mitotic Timing Analysis

RIT1^M90I^-expressing cells were generated by transduction with a pLX303-RIT1^M90I^ lentivirus for and selected with puromycin. One day before imaging, cells were seeded at a density of 30,000 cells per well of an 8-well Ibidi glass-bottomed plate. For drug-treated populations, 0.5uM Reversine (Selleckchem) was added two hours before imaging. Live-cell imaging was performed using a 20×/0.70 Plan Apo Leica objective on an automated Leica DMi8 microscope outfitted with an Andor CSU spinning disk unit equipped with Borealis illumination, an ASI automated stage with Piezo Z-axis top plate, and an Okolab controlled environment chamber (humidified at 37°C with 5% CO2). Long term automated imaging was driven by MetaMorph software (version 7.10.0.119). Images were captured with an Andor iXon Ultra 888 EMCCD camera. Images were captured every minute for 18 hours. Time in mitosis was measured as time from nuclear envelope breakdown to the onset of anaphase. Imaging experiment was repeated four times with distinct biological replicates and 50 cells were analyzed per cell line per condition.

Mitotic index was calculated from one time point selected from live-image experiments in HeLa H2B parental and HeLa H2B RIT1^M90I^ described above. At each time-point, three independent, representative images were collected and the mitotic index, the number of cells undergoing mitosis (in any phase between prometaphase to telophase) divided by the total number of cells was calculated. The same time point was analyzed in two independent experiments for a total of 6 total mitotic index analyses per cell line.

### Functional enrichment analysis

Gene set enrichment analysis was performed using the MsigDB ^75 76^ and StringDB ^77^ databases. For all analyses an FDR cut-off of < .05 was used. To account for differences in the number of cooperating factors between RIT1^M90I^ and KRAS^G12V^, overlap analysis was carried out on the top 152 positively-selected genes from each screen.

### TCGA RNA expression analysis

*RIT1* RNA-seq data from TCGA lung adenocarcinomas (230 samples)^12^ was obtained from cBioPortal for Cancer Genomics (http://cbioportal.org). The z-score for *RIT1* mRNA expression is determined for each sample by comparing *RIT1* mRNA expression to the distribution in a reference population harboring typical expression for the gene^78^.

For differential gene expression analysis, *RIT1*-altered samples were defined as tumors with either mutation of *RIT1* (n=5) or amplification of *RIT1* (n=32). The remaining 193 tumors were considered non-*RIT1*-altered. Differential expression analysis was performed in cBioPortal, and genes with q-value <= 0.25 were downloaded from cBioportal (http://cbioportal.org). Pathway enrichment analysis was carried out on the 1015 under-expressed genes identified in *RIT1-*altered sample using the StringDB ^77^ database with an FDR < .05.

For analysis of individual Hippo pathway genes in *RIT1* altered samples, we included the following genes as Hippo pathway genes: *AMOTL1*, *NF2*, *STK4*, *STK38L MAP4K5*, *TAOK3*, *SAV1*. To determine the portion of *RIT1* altered tumors with low expression of any one Hippo pathway gene, low expression was defined as >1 standard deviation lower than the mean expression of the 230 sample dataset. Then we determined the proportion of samples with low expression of at least one of the above genes, or with normal expression of all genes.

## STATISTICAL ANALYSIS

### Genome-wide CRISPR screen

A log_2_-normalized count matrix of sequencing reads mapped to sgRNAs was used as an input to MAGeCK ^32^. sgRNAs with <1 read-per-million in the sequencing of the Brunello library plasmid were excluded from the dataset. The following sgRNAs were determined to not target the introduced cDNA oncogenes, and so were excluded from the dataset:

KRAS: (AGATATTCACCATTATAGGT)
RIT1: (CATGCGGGACCAGTATATGA,GTGATGATCTGGCTTACCAA)
EGFR: (TGTCACCACATAATTACCTG)

Reproducibility of the screen was assessed by calculating Pearson correlations of pairwise replicate comparisons for each replicate set (n = 2-3 replicates per screen arm). Pearson correlations ranged from 0.58-0.94 (median = 0.82). Scores from early time-point (ETP) samples were highly correlated to plasmid DNA (median = 0.93; **Supplementary Fig. 2d**) so comparison to plasmid DNA was used for all subsequent analyses. Guide RNAs were collapsed to gene scores using MAGeCK, and log_2_ (fold-change) values (LFC) were computed compared to starting guide RNA abundance in plasmid DNA. To check the performance of each screen, we calculated robust strictly standardized mean difference (SSMD) statistics^6^ for each replicate, comparing the LFC between non-targeting sgRNAs and LFC values of sgRNAs targeting the spliceosomal, ribosomal, and proteasomal gene sets from KEGG ^79^. All endpoint replicates passed the quality control threshold of SSMD < −0.5, with the median SSMD = −3.9 (**Supplementary Table 1**). Normalization across screens was performed using methods similar to those previously described^6^. First, the LFC data were normalized within each replicate by subtracting the median LFC of the replicate and then dividing by the median average deviation. Next, we scaled each replicate based on the LFC of pan-essential genes and pan-non-essential genes using previously defined lists by DepMap^70^. The data were scaled such that the median of the non-essential genes in each replicate is 0 and the median of the essential genes is −1. “CRISPR scores” were defined as this scaled, normalized LFC data. The final normalized, scaled data are supplied as (**Supplementary File 2**). For analysis of expressed vs. non-expressed genes (**Supplementary Fig. 2f**), we defined expressed genes from RNA-sequencing data of PC9 cells^80^ as genes with log_2_ (transcripts per million - TPM) > 2 and non-expressed genes as log_2_TPM<1.

### Statistical Tests

Statistical tests are indicated in the figure legends. Results were analyzed for statistical significance with GraphPad Prism 8 software or R (v 3.6.3). A *p* value of < 0.05 was considered significant (\**p* < 0.05, \*\**p* < 0.001, \*\*\**p* < 0.001, \*\*\*\**p* < 0.001).

## Data Availability Statement

RNA-seq data generated by TCGA were downloaded from cBioPortal for Cancer Genomics (http://cbioportal.org). The authors declare that all other data supporting the findings of this study are available within the paper and its Supplementary information files.

## Acknowledgements

We thank Amy Goodale (Broad Institute) for technical assistance in the design and synthesis of the targeted validation screening library and Dr. Daphne Avgousti (Fred Hutchinson Cancer Research Center) for providing HeLa H2B-GFP cells. We thank the Broad Institute Genetic Perturbation Platform for technical assistance and advice and the Fred Hutch Genomics Shared Resource (supported by NIH/NCI Cancer Center Support Grant P30 CA015704). This research was funded in part through NCI grant R00CA197762 to A.H.B, donations from the Smith family to A.H.B., NIH/NCI Cancer Center Support Grant P30 CA015704 New Investigator support to A.H.B., a Lung Cancer Research Foundation Research Grant to A.V., the Hutch United postdoctoral fellowship to A.V.. A.R. was supported in part by PHS NRSA T32GM007270 from NIGMS. P.C.R.P. was supported in part by NSF DGE-1762114.

## Contributions

A.H.B. conceived of and supervised the project. A.V., F.P., E.H., and A.H.B. designed experiments. A.V., N.T.N., A.R., P.C.R.P., and A.H.B. analysed the data. A.V., N.N., A.R., P.C.R.P., J.K.L., F.D., J.C., J.W., M.R., and A.H.B., conducted experiments. A.V. and A.H.B. wrote the manuscript. All the authors critically reviewed the manuscript and approved the final version.

## SUPPLEMENTARY FIGURE LEGENDS

**Supplementary Fig. 1. a**, Western blot of lysates from SALE and AALE cells stably expressing wild-type or mutant *RIT1* or mutant *RAS* constructs, and empty vector (control). Vinculin and total MEK were used as loading controls. p-MEK, phosphorylated Ser217/221. **b**, Schematic of the 135bp deletion identified by Sanger sequencing and ICE analysis in *RIT1* knockout clones generated at the *RIT1* locus by transfection with three synthetic sgRNAs. **c**, Western blot of RIT1 expression in clonal *RIT1* knockout cells compared to parental PC9-Cas9-RIT1^M90I^ cells or control PC9-Cas9-luciferase cells. **d**, 96 hour dose-response curve of clonal *RIT1* knockout in osimertinib. Viable cell fraction was determined by CellTiterGlo luminescence of treated vs. DMSO-treated cells. Data shown is the mean + s.d. of two technical replicates.

**Supplementary Fig. 2. a**, Western blot of lysates generated from parental PC9 cells or stable PC9 cell pools expressing Cas9 and indicated oncogenes. Actin served as a loading control. The primary antibodies used are indicated on the right. **b**, Distribution of fluorescence in stable Cas9-expressing PC9 cells transduced by EGFP-targeting sgRNAs (red), or non-Cas9 expressing PC9 cells transduced by EGFP-targeting sgRNAs (blue), and non-fluorescent PC9 cells (gray). **c**, Schematic illustrates the workflow of genome-wide CRISPR/Cas9 knockout library screening. On day 1 the human genome-wide Brunello CRISPR/Cas9 knockout library (Brunello) was transduced into isogenic PC9 cells at low MOI. The transduced cells were selected by puromycin on day 2. Cells were cultured in vehicle (DMSO) or erlotinib for ~12 population doublings. Genomic DNA was extracted from erlotinib and vehicle treated cells and the sgRNA sequence was amplified by PCR. Abundance of sgRNAs was determined by Illumina sequencing and analyzed by MAGeCK^32^. **d**, Pairwise Pearson correlation matrix of sgRNA abundance across screen replicates. Each column/row corresponds to a different replicate. **e**, Volcano plot showing the distribution of targeting guides, essential guides, and control non-targeting guides (collapsed at gene level and/or average of four sgRNA per gene) after normalization. “CRISPR Score” was calculated as described in the Method. Data shown is from PC9-Cas9-RIT1^M90I^ but similar results were obtained from each cell line. **f**, log_2_ fold-change distribution of all sgRNAs targeting expressed genes (red) or non-expressed genes (blue) in each cell line. Vertical lines indicate the mean of each distribution. **g-i**, Expanded view of data shown in Fig. 2d. Box plot showing the log_2_sgRNA abundance (reads per million) from sgRNAs targeting **g**, *KRAS*, **h**, *RIT1* or **i**, *EGFR* in isogenic PC9-Cas9 cells by screening condition. Plasmid, starting plasmid library. Control, Renilla luciferase vector control cells. ETP, early time point. DMSO, vehicle control. Box plots show the median (center line), first and third quartiles (box edges), and the min and max range (whiskers) of replicates. *** *p* < 0.001, calculated by MAGeCK (methods). For control PC9 cells ETP and DMSO n= 3 biological replicates. For oncogene-expressing PC9 cells DMSO and erlotinib n= 2 biological replicates.

**Supplementary Fig. 3. a-f**, Box plot showing the log_2_sgRNA abundance (reads per million) from sgRNAs targeting key genes discussed in the text. Labeling is as in Supplementary Fig. 2g-i. * *p* < 0.05, ** *p* < 0.01, *** *p* < 0.001 calculated by MAGeCK (Methods). For control PC9 cells ETP and DMSO n= 3 biological replicates. For oncogene-expressing PC9 cells DMSO and erlotinib n= 2 biological replicates. **g**, Primary and secondary CRISPR screen correlation analysis in PC9-Cas9-RIT1^M90I^ erlotinib-treated cells. The solid black diagonal line displays the linear regression. Relevant RIT1^M90I^ dependencies and cooperating genes discussed in the text are labeled in red.

**Supplementary Fig. 4. a**, Rank plot of the mean difference in CRISPR score of PC9-Cas9-RIT1^M90I^ erlotinib-treated and vehicle (DMSO) treated cells of secondary validation screen. Selected essential genes are labeled in red. **b**, Validation of *USP9X*, *AURKA*, and *SHOC2* dependencies. Data shown is 8 sgRNAs across three biological replicates of each condition. Labeling is an in Supplementary Fig. 2g-i. * *p* < 0.05, *** *p* < 0.001, calculated by unpaired two-tailed t-test. **c**, Schematic of the deletions identified by Sanger-sequencing and ICE analysis in *SHOC2*, *USP9X*, and *AURKA* knockout clones generated by transfection with synthetic multi-guide sgRNA guides. **d**, Western blot of lysates generated from PC9-Cas9-RIT1^M90I^ cells or clonal PC9-Cas9-RIT1^M90I^ + *USP9X* knockout (KO) cells. Vinculin was used as a loading control. **e**, 96 hour dose-response curve of osimertinib in clonal *SHOC2* knockout (KO), *USP9X* KO, or *AURKA* KO cells derived from PC9-Cas9-RIT1^M90I^ cells. The same data for control and PC9-Cas9-RIT1^M90I^ cells is plotted on each panel for reference. Data shown is the mean ± s.d. of 2 technical replicates. **f**, Area under the curve (AUC) analysis of data from panel (e).

**Supplementary Fig. 5. a-f**, Box plot showing the log_2_sgRNA abundance (reads per million) from sgRNAs targeting key Hippo pathway genes across isogenic PC9-Cas9 cells line by condition. Labeling is as in Supplementary Fig. 2g-i. * *p* < 0.05, ** *p* < 0.01, *** *p* < 0.001, calculated by MAGeCK. **g**, Validation of *NF2* positive selection from the secondary validation screen in PC9-Cas9-RIT1^M90I^ cells. Data shown is 8 sgRNAs across three biological replicates of each condition. Box plots show the median (center line), first and third quartiles (box edges), and the min and max range (whiskers) of replicates, *** *p* < 0.001 by unpaired two-tailed t-test. **h**, Western blot of lysates from PC9-Cas9-RIT1^M90I^ cells or stable PC9-Cas9-RIT1^M90I^ + *NF2* knockout (KO) cell pools. Vinculin was used as a loading control. **i.** Dose-response curve of erlotinib in PC9-Cas9-RIT1^M90I^ + *NF2* knockout cell pools. Cells were treated with erlotinib for 96 hours. Viable cell fraction was determined by CellTiterGlo luminescence of treated cells normalized to the average value of DMSO-treated cells. Data shown is the mean + s.d. of 2 technical replicates. **j**, Frequency of *RIT1* amplification and mutation (n=37) from the TCGA lung adenocarcinoma marker paper^12^ visualized by cBioportal (http://cbio.mskcc.org). Each column represents a sample. Samples lacking alterations (n=193) in *RIT1* are not shown. 16% indicates the percentage of samples in the cohort with mutation or amplification in *RIT1* (37/230). **k**, Box plot showing the relationship between *RIT1* mRNA abundance and copy number of *RIT1* from TCGA analysis shown in (j). Diploid, two alleles present; Gain, low-level gene amplification event; Amplification, high-level gene amplification event. *** *p* < 0.001 by unpaired two-tailed t-test.

