## Supplementary Figures for "An integrative oncogene-dependency map identifies unique vulnerabilities of oncogenic EGFR, KRAS, and RIT1 in lung cancer"

### Supplementary fig. 1

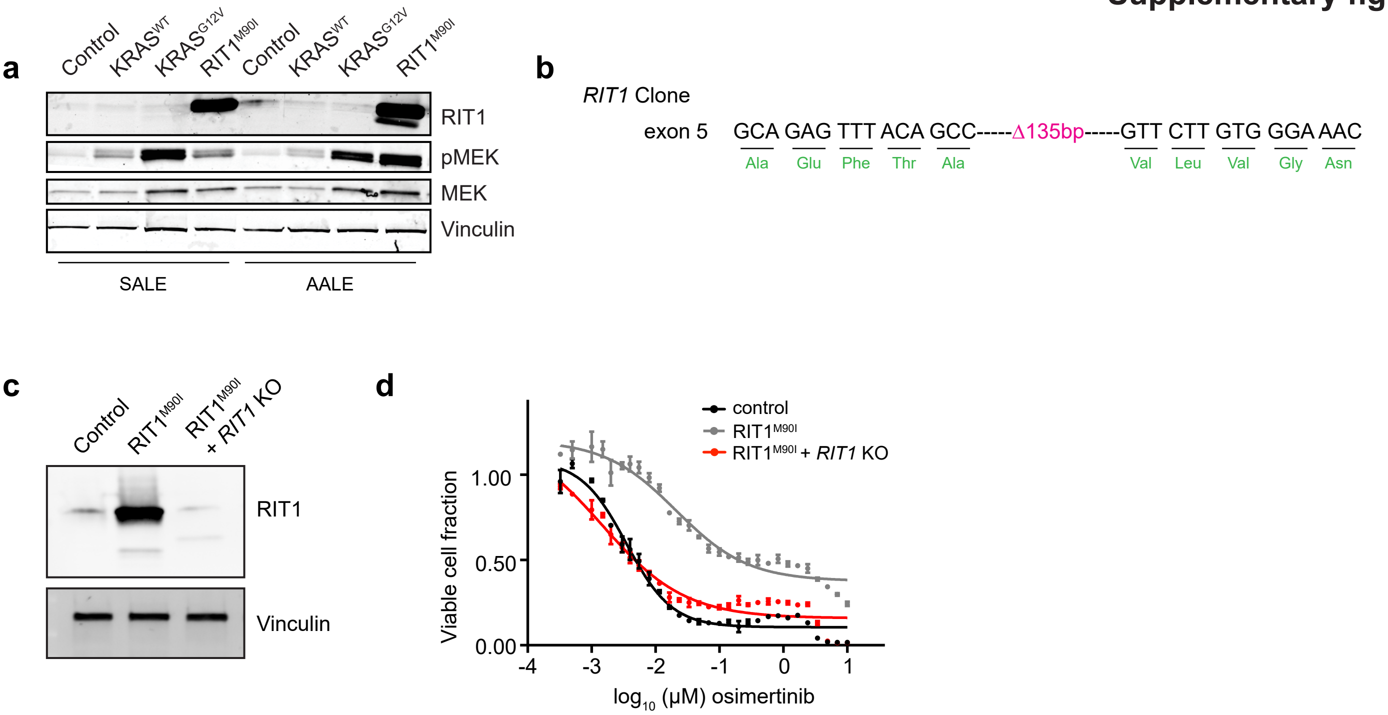

### Supplementary fig. 2

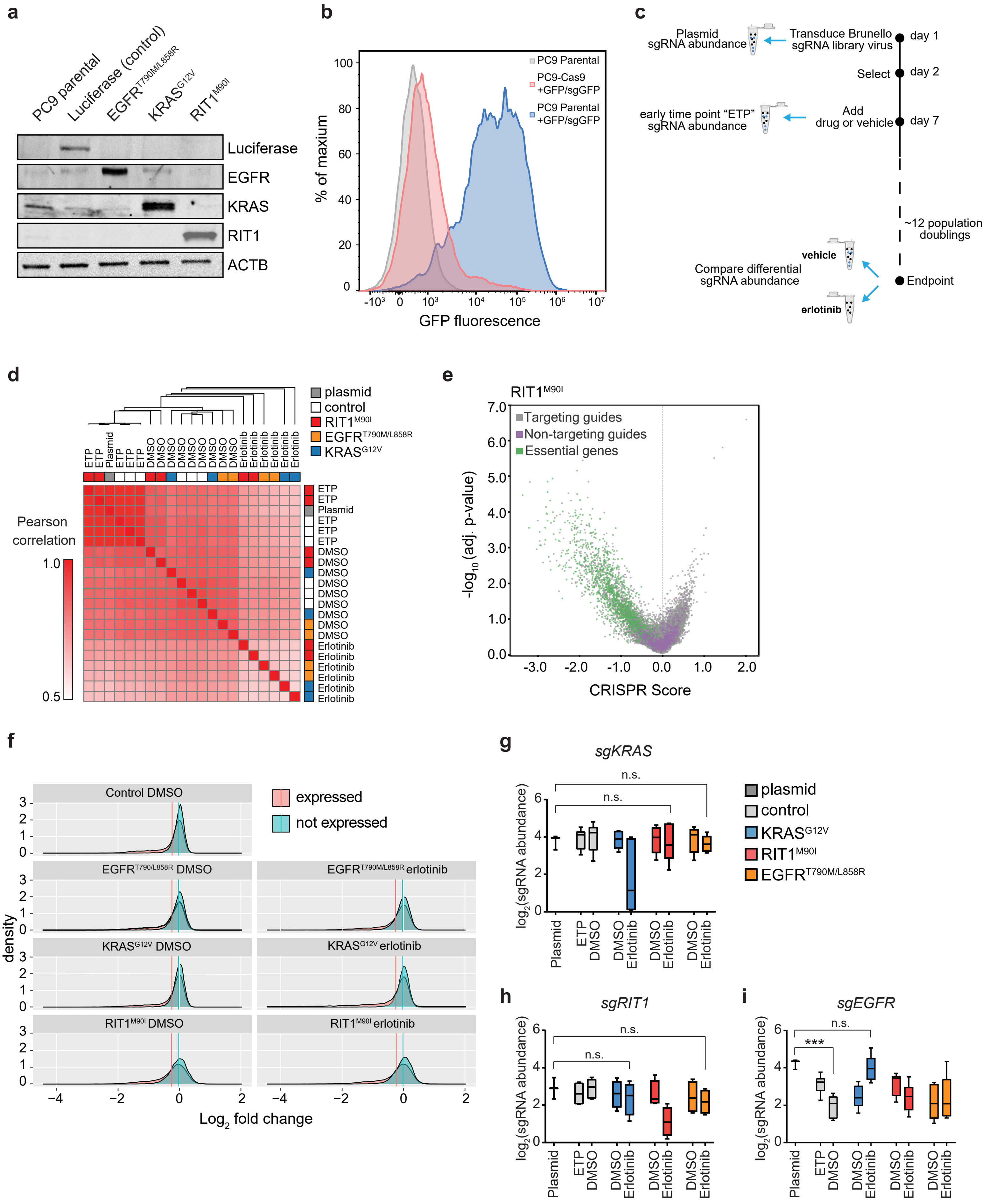

### Supplementary fig. 3

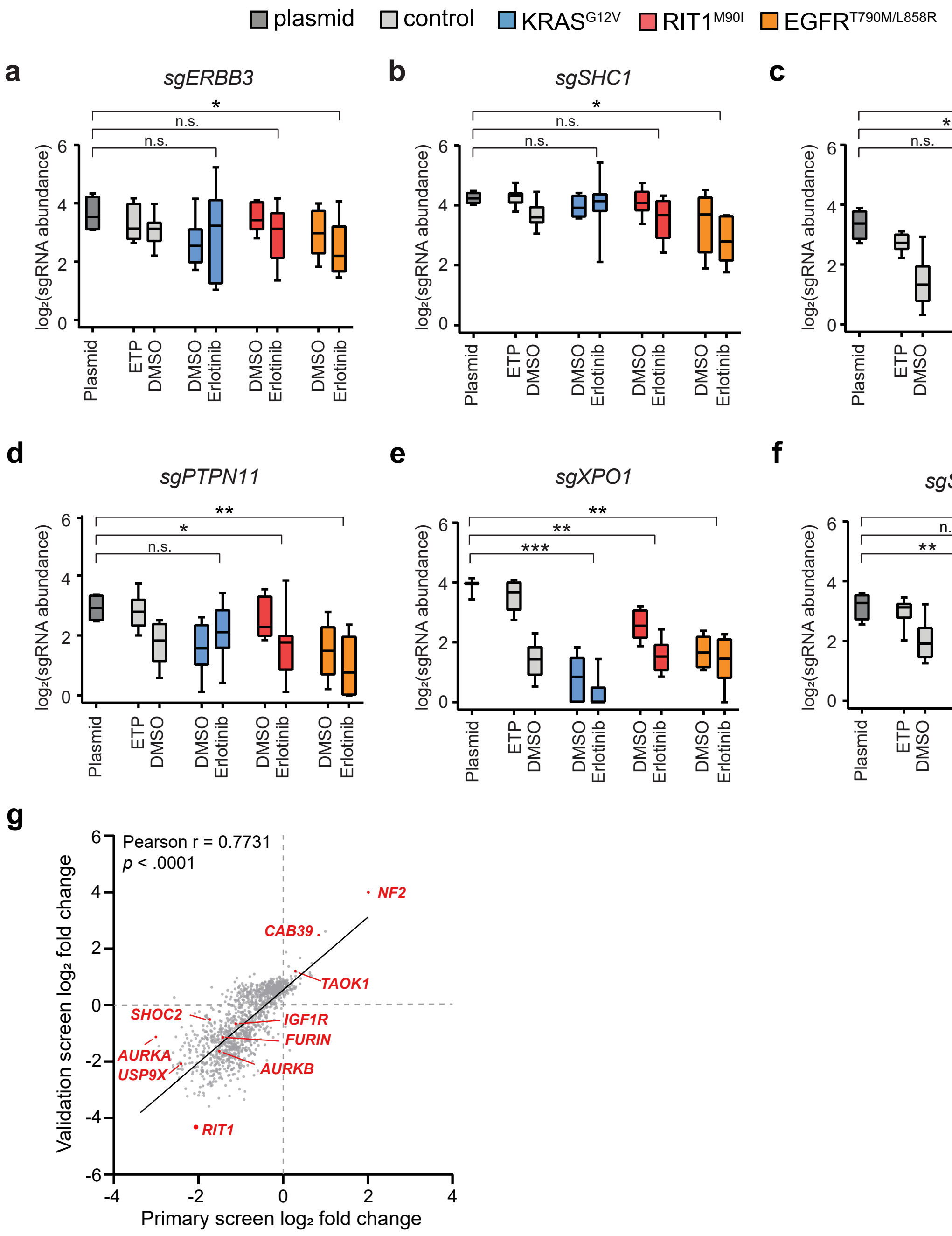

**Supplementary fig. 4**

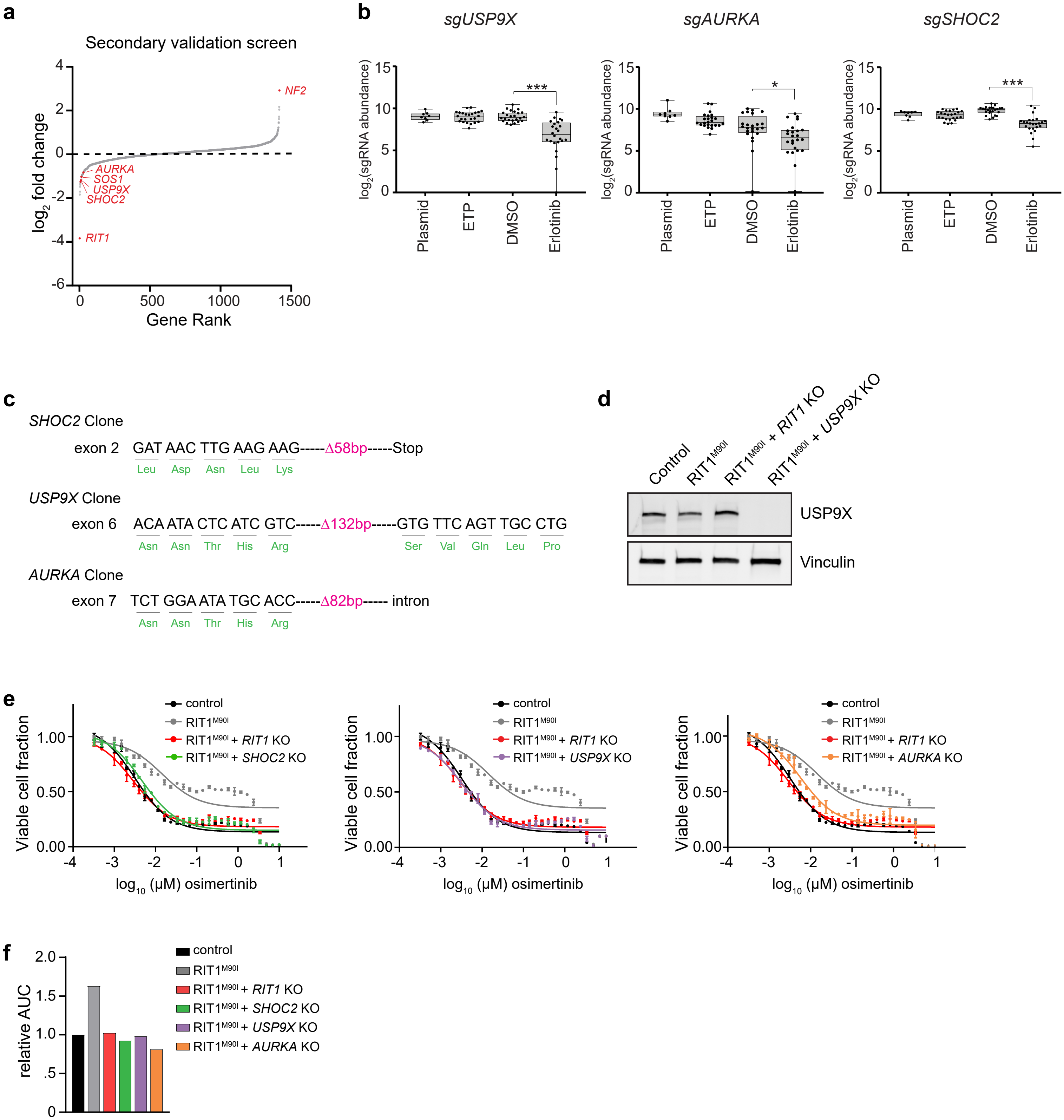

### Supplementary fig. 5

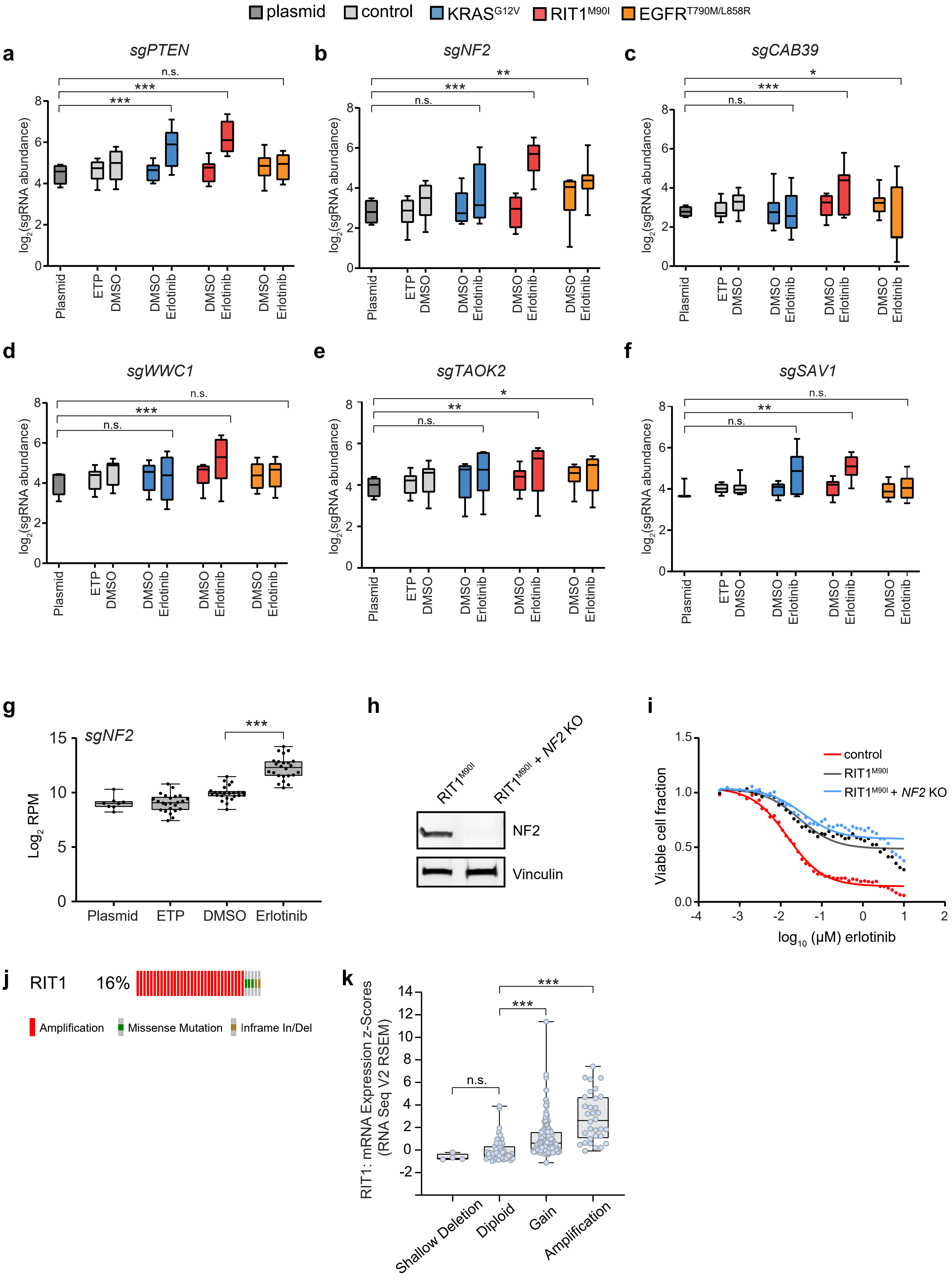
